# Analysis of gliomas DNA methylation: Assessment of pre-analytical variables

**DOI:** 10.1101/2024.03.26.586350

**Authors:** Karol Bomsztyk, Daniel Mar, Oleg Denisenko, Suzanne Powell, Monika Vishnoi, Jennifer Delegard, Anoop Patel, Richard G Ellenbogen, Rohan Ramakrishna, Robert Rostomily

## Abstract

Precision oncology is driven by molecular biomarkers. For glioblastoma multiforme (GBM), the most common malignant adult primary brain tumor, O6-methylguanine-DNA methyltransferase (*MGMT*) gene DNA promoter methylation is an important prognostic and treatment clinical biomarker. Time consuming pre-analytical steps such as biospecimen storage before fixing, sampling, and processing are major sources of errors and batch effects, that are further confounded by intra-tumor heterogeneity of *MGMT* promoter methylation. To assess the effect of pre-analytical variables on GBM DNA methylation, tissue storage/sampling (CryoGrid), sample preparation multi-sonicator (PIXUL) and 5-methylcytosine (5mC) DNA immunoprecipitation (Matrix MeDIP-qPCR/seq) platforms were used. *MGMT* promoter CpG methylation was examined in 173 surgical samples from 90 individuals, 50 of these were used for intra-tumor heterogeneity studies. *MGMT* promoter methylation levels in paired frozen and formalin fixed paraffin embedded (FFPE) samples were very close, confirming suitability of FFPE for *MGMT* promoter methylation analysis in clinical settings. Matrix MeDIP-qPCR yielded similar results to methylation specific PCR (MS-PCR). Warm ex-vivo ischemia (37°C up to 4hrs) and 3 cycles of repeated sample thawing and freezing did not alter 5mC levels at *MGMT* promoter, exon and upstream enhancer regions, demonstrating the resistance of DNA methylation to the most common variations in sample processing conditions that might be encountered in research and clinical settings. 20-30% of specimens exhibited intratumor heterogeneity in the *MGMT* DNA promoter methylation. Collectively these data demonstrate that variations in sample fixation, ischemia duration and temperature, and DNA methylation assay technique do not have significant impact on assessment of *MGMT* promoter methylation status. However, intratumor methylation heterogeneity underscores the need for histologic verification and value of multiple biopsies at different GBM geographic tumor sites in assessment of *MGMT* promoter methylation. Matrix-MeDIP-seq analysis revealed that *MGMT* promoter methylation status clustered with other differentially methylated genomic loci (e.g. HOXA and lncRNAs), that are likewise resilient to variation in above post-resection pre-analytical conditions. These *MGMT*-associated global DNA methylation patterns offer new opportunities to validate more granular data-based epigenetic GBM clinical biomarkers where the CryoGrid-PIXUL-Matrix toolbox could prove to be useful.

## BACKGROUND

Glioblastoma (GBM) is the most common adult-type diffuse glioma, with a universally poor prognosis (1). The application of molecular biomarkers have a significant impact on the histological typing and diagnosis of GBM, as well as predicting patient survival and response to treatment (2–4). For IDH WT GBM, the methylation status of the O^6-^methylguanine-DNA methyl-transferase (*MGMT)* gene promoter is the only molecular biomarker currently utilized as a clinical assay to stratify pateint outcomes and responses after standard of care temozolomide (TMZ) and radiation therapy (4–10). However, outlying cases that show a lack of correlation of the promoter methylation with clinical outcomes suggest that unrecognized effects of pre-analytic variables in tissue sampling, processing and assay techniques may contribute to these discrepancies (11,12).

Advances in biotechnologies and an increased understanding of tumorigenesis have provided unprecedented opportunities for the discovery of cancer biomarkers and their applications in clinical oncology (13). While the cancer biomarker field has historically been focused on developing and improving molecular assays, relatively fewer research efforts have been made to find ways to maximize the quality of biospecimens and assess the impact of pre-analytic variables before clinical testing (14). As such, despite its pivotal clinical importance, there has been little systemic analysis of how pre-analytic variables such as fixation, warm and cold ischemia times and sampling of a heterogenous tumor may impact measurement of *MGMT* promoter methylation. Moreover, our understanding of the potential mechanistic or prognostic significance of *MGMT* promoter methylation as a reporter of multiome-wide alterations is incomplete.

Every biomarker analysis begins with biospecimen collection, sampling, and analyte extraction which are critical variables in determining data reproducibility (15). Sample preparation, often involving multiple steps, remains the most challenging and time-consuming component of biomarker analysis and a principal source of data irreproducibility (16,17). Consistent with this notion, most laboratory medicine errors are estimated to occur at the pre-analytical phase (17) and incomplete understanding of the impact of pre-analytical procedures therefore may account for irreproducibility of some biomarker assays (14,18–22). As a consequence, false positive or negative biomarker assays resulting from insufficiently tested biospecimen protocols can confound drug development, diagnostic tests, cancer treatments and represent a critical impediment to advances in precision medicine. In addition, institution or clinician-specific sample acquisition and processing factors may also impact assay reproducibility (14,21). Therefore, to achieve maximal clinical value from biomarker assays, standard pre-analytical operating procedures are needed to ensure that individuals’ resected tumors are maximally preserved for molecular assays to faithfully report native cellular and molecular signatures (14,21).

There are >2500 publications in PubMed listed under *MGMT* promoter methylation, but none appear to have examined pre-analytical variables that might influence outcomes of this biomarker assay. To address this deficiency the current study systematically measured the impacts of pre-analytic factors on *MGMT* promoter and other methylated loci including time of *ex-vivo* ischemia, fixation/storage, sampling methods, and intratumor heterogeneity.

## MATERIALS, DEVICES, BIOSPECIMENS, AND METHODS

Details of each section listed below are provided in the Supplement.

### Materials

Buffer recipes are listed in the Supplementary section. Kit/enzyme/antibodies catalog numbers commercial suppliers and PCR primers also listed in the supplementary tables **(Table S1-3).**

### Devices

The CryoGrid system (23) was used to cryostore (CryoTray) and sample (CryoGrid) tissue, and PIXUL (24) multi-sonicator was used to extract and prepare DNA and RNA.

### Biospecimens

Glioma specimen collection was done according to protocols approved by the Institutional Ethical Review Boards at the Houston Methodist Research Institute, Houston, TX and at the University of Washington, Seattle, WA. Cell cultures were derived from GBM patient specimen. Xenograft model was generated using U87-MG cell lines injected subcutaneously at the flank of nude mice (25). All animal studies were performed according to protocols approved by the Institutional Animal care and Use Committee (IACUC) at the Houston Methodist Research Institute (see more details in supplementary material).

### Methods

#### Frozen and FFPE tissues

Tissues were stored in CryoTrays at -80°C (Frozen) (23). FFPE blocks were prepared from matched frozen tissues as described in (26) and sectioned using a microtome. Prior to analyte extractions, FFPE curls were deparaffinized and rehydrated (27).

#### Analysis of methylated DNA

Methylated DNA immunoprecipitation in 96-well microplates (Matrix-MeDIP) (28) was followed by either qPCR/PCRCrunch analysis of selected loci or library production, sequencing, and generation and analysis of BAM and bigwig files (29). Methylation specific PCR (MS-PCR) was done as described previously described (30).

#### Analysis of mRNA

Reverse transcription was done in 96-well microplates (Matrix-RT) (23). cDNAs were analyzed by qPCR/PCRCrunch (31) .

#### H&E slide preparation and digital image analysis

Flash frozen tissue fragments immobilized in CryoTrays were used to extract two-three 1x2 mm cores using the CryoCore. Cores were formalin-fixed and processed via paraffin embedding on an automated tissue processor. Tissue cores embedded in paraffin were sectioned at 4-5 μm onto charged slides, baked at 60^0^C, stained with routine H&E, and cover-slipped with permanent mounting media. Whole slide digital imaging was performed on an Aperio AT Turbo or Aperio Versa 200 at 20X.

## RESULTS AND DISCUSSION

### Methylated DNA immunoprecipitation (MeDIP) to assess *MGMT* promoter methylation (**Fig.1**)

The *MGMT* promoter-exon 1 region (-953 to +202 bp relative to transcription start site) contains 97 CpG sites, within a 777bp GC island, that can accept methyl groups and as a result can contribute to repressing transcription (32,33). *MGMT* promoter methylation studies have most commonly been done using either methylation-specific PCR (MS-PCR) or pyrosequencing (5,9). Both methods require bisulfite conversion (34), as do several other *MGMT* methylation assays (35,36). Newer DNA methylation methods are being introduced that are bisulfite-free (37,38). MS-PCR remains the main *MGMT* methylation analysis method used in clinical laboratories (36). This is a single site-based methylation assay. Methylation status of several *MGMT* promoter CpGs have been shown to help predict response to therapies (39). Most of these *MGMT* promoter methylation single-site(s) analysis methods are semi-quanatitative rather than quantitive. Further, there is no convincing evidence that any specific methylated CpG *MGMT* promoter sites are better clinical predictors than others (36). Instead, integrated cluster set of methylated CpGs not only at the *MGMT* promoter but also genome-wide, along with other assays (e.g., mRNA and protein expression), are emerging as better cancer clinical predictors (e.g. *HOXA*) (40,41) .

**Fig. 1.**
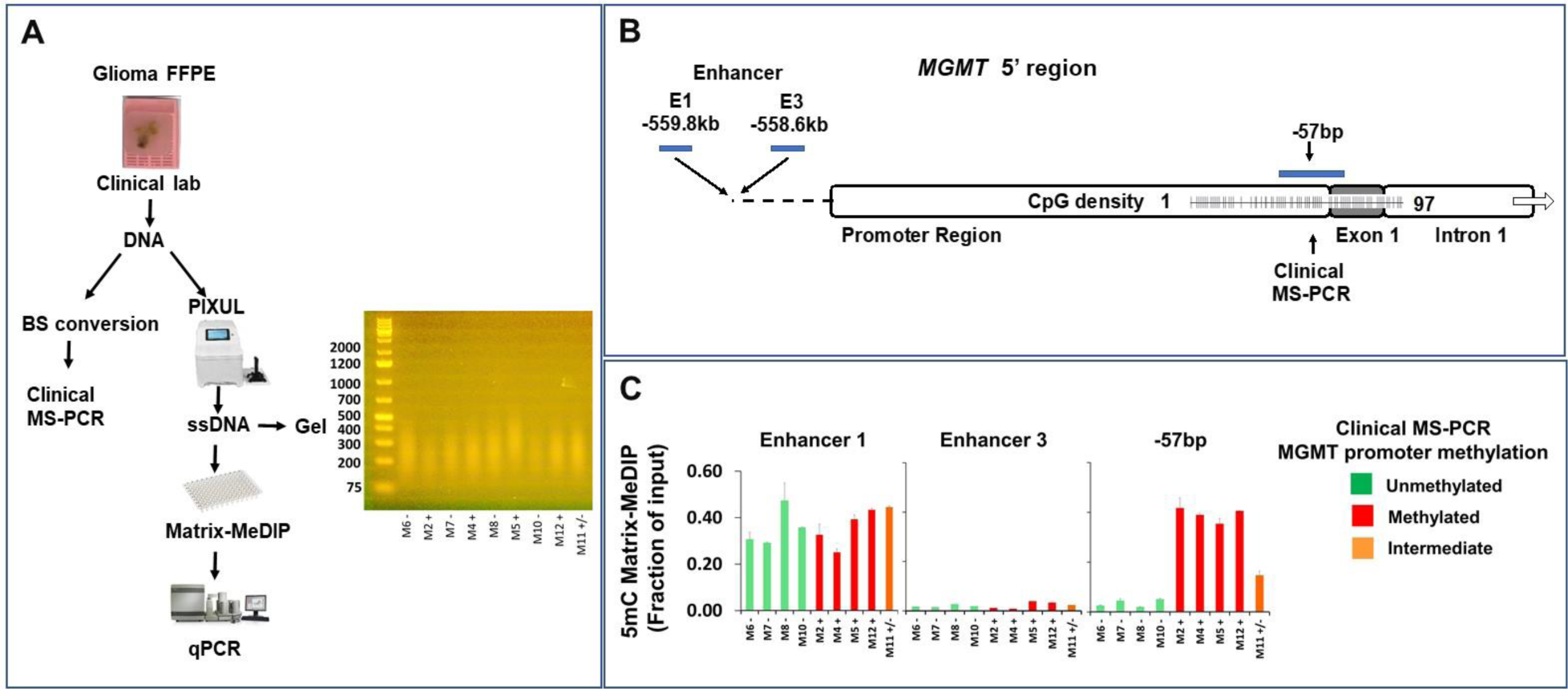
Matrix-MeDIP-qPCR *MGMT* promoter methylation analysis of clinical laboratory GBM DNA samples. ***A.*** DNA was isolated from FFPEs by UW Tumor Clinical Laboratory per their protocol. The DNA was either analyzed by bisulfite conversion (BS) followed by methylation-specific PCR, MS-PCR or fragmented in 96-well plated in PIXUL device, boiled to obtain ssDNA, and assayed in 96-well Matrix-MeDIP using 5mC antibody followed by qPCR. ***B.*** Shows the structure of the MGMT gene 5’-regions with location of PCR primers used. ***C.*** Matrix-MeDIP-qPCR results. Colors show UW Clinical MS-PCR results reported as either *MGMT* promoter methylation negative, positive, or intermediate MGMT promoter methylation, Mean ±SEM, n=4 qPCR.

We have previously developed a 96-well microplate-based high throughput methylated DNA immunoprecipitation assay, Matrix MeDIP, that does not use bisulfite conversion and uses sheared DNA (28,29,42). MeDIP is a quantitative assay which averages CpG methylation levels over sheared DNA fragment length. Applied to DNA methylation analysis of *MGMT* promoter region (32), MeDIP could be useful diagnostically to improve patient stratification. Further, MeDIP allows to readily assay selected sites (qPCR) or carry out genome-wide (NGS) analysis (29). To validate Matrix-MeDIP for *MGMT* methylation analysis we designed PCR primers within the promoter CpGs and carried out a series of experiments to compare Matrix-MeDIP with MS-PCR. We obtained anonymized DNA samples from the University of Washington Clinical Diagnostic laboratory that were previously tested for *MGMT* promoter methylation by MS-PCR, a total of 9 samples: 4 unmethylated, 4 methylated, and 1 intermediate **(Fig.1).** DNA was sheared using PIXUL (24) and boiled, and the ssDNA was immunocaptured using anti-5mC antibody in Matrix MeDIP. The average size of the sheared DNA was 200-250bp **(Fig.1A).** qPCR primers were used to the *MGMT* CpG promoter region that include the site used in the clinical MS-PCR assay (-126/+12bp) as well as two regions in the 5’ intergenic *MGMT* enhancer (43), **(Fig.1B).** As shown in **Fig.1C**, at the -126bp to +12bp region (overlaping with -57 bp site used in clinical MS-PCR assays), Matrix MeDIP revealed DNA methylation levels that match methylated vs. unmethylated *MGMT* promoter status defined by the MS-PCR clinical diagnostic assay at the single site (-57bp). By contrast, there were no differences in methylation at 5’-intergenic enhancer sites in the same assay. Consistent with the higher number of CpGs the more upstream enhancer (E1) **(Fig.S1)** was highly methylated while the more downstream enhancer site (E3) with only a couple of CpGs showed little or no 5mC signal **(Fig.1C).** Note that the sheared DNA fragments are 100-150bp longer than the amplicon, allowing for the detection of methylated CpGs sites flanking the amplicon regions **(Fig.S1).**

To further validate MeDIP, PIXUL-based workflow was used to isolate DNA from GBM-derived cell cultures from tumors with methylated and unmethylated *MGMT* promoter. **Fig.2** shows that 5mC levels at two regions of the *MGMT* promoter bearing CpGs **(Fig.S1)**, designated as -363bp and -57bp relative to the TSS, were high in GBM-derived cultures from individuals with methylated promoter and very low in those individuals with unmethylated promoter. At the enhancer 1, the 5mC levels were high but not different across the GBM-derived cultures. The 5mC signal was low across all samples at the promoter -903bp and enhancer 3 sites which have only couple of CpGs **(Fig.S1).** These data further validate the PIXUL-Matrix-MeDIP-qPCR method for assessment of *MGMT* promoter DNA methylation.

**Fig. 2.**
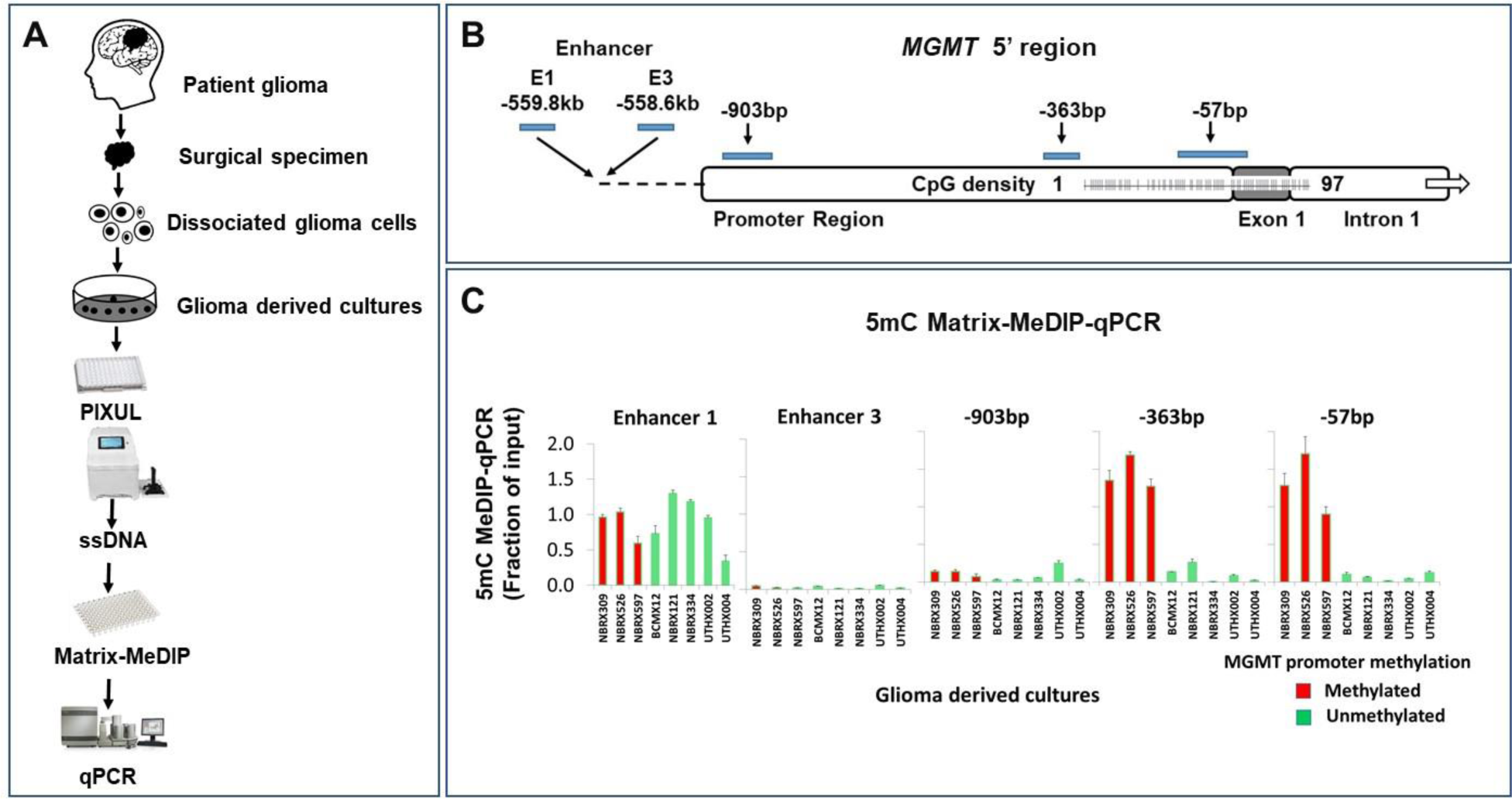
Matrix-MeDIP-qPCR DNA methylation analysis of GBM-derived cultures. ***A.*** 96-well plate using PIXUL sonicator was used to isolate DNA from cultures derived from MGMT promoter methylated (positive, *red*) and unmethylated (negative, *green*) GBMs. After boiling, ssDNA was assayed in 96-well plate Matrix-MeDIP-qPCR using 5mC antibody. ***B.*** MGMT 5’ region showing sites of PCR primers. ***C.*** Matrix-MeDIP-qPCR results, Mean ±SEM, n=3.

### Distribution of *MGMT* promoter and enhancer DNA methylation levels across a large surgical collection of paired frozen and FFPE tissue fragments

We have assessed DNA methylation levels at the *MGMT -*57bp promoter and enhancer 1 regions across 173 surgical previoulsy flash-frozen specimens, from 90 individuals (histology as follows: 76 GBM, 3 high grade gliomas, 4 astrocytomas, 1 oligodendroglioma, 5 unknown, 1 indetermined). Frozen tissues in CryoTrays were sampled with CryoCore, cores were jetted into 96-well microplates, DNA was isolated using PIXUL, and ssDNA was assessed in Matrix MeDIP-qPCR **(Fig.S2A-B).** The distribution of methylation levels, determined by Matrix-MeDIP, across all surgical biospecimens is quite different at the promoter and enhancer sites **(Fig.S2C-D).** Heterolon is a data metric that we developed to quantify inter-tumor and intra-tumor heterogeneity - the greater the heterolon, the greater the variation (Methods, supplement). The heterolon at the -57bp promoter site is higher, 1.09, than it is at the enhancer 1, 0.55 **(Fig. S3**), indicating greater variation at the promoter site. In clinical practice and studies, gliomas are qualitatively stratified into *MGMT* promoter unmethylated, intermediate, or methylated tiers (39,44). Based on the dataset distribution we devised a three-tier stratification of samples based on Matrix-MeDIP-measured *MGMT* promoter methylation levels in tumor fragments as follows: i) below median as unmethylated, ii) between median and mean as intermediate, and iii) above mean as methylated. We applied the above overall approach for biospecimen MeDIP-qPCR dataset analysis in evaluating contribution of pre-analytical variables.

Formalin fixation and paraffin embedding (FFPE) is the most commonly used method for tissue specimen storage. Importantly, FFPE blocks are often used in clinical settings to transport GBM samples to clinical labs for *MGMT* promoter methylation analysis (10). FFPE tissues are also widely used for longitudinal specimen storage (45,46). This practice has generated vast tissue repositories, including GBMs (45). Further, improvements in FFPE samples processing are demonstrating superior practicality of FFPE over frozen samples for molecular assays in research and clinical settings (29,47,48). FFPE specimens are stored at room temperature, are easy to transport, and pose less of a biohazard than non-fixed tissues. Formaldehyde cross-links amino groups with nearby (∼2Å) nitrogen atoms in proteins, DNA, or RNA. Formaldehyde cross-linking can be reversed, which allows these blocks to be exploited for molecular analysis including DNA methylation (29,49,50). And yet, formalin may stimulate 5mC deamination, an important pre-analytical modification than may alter results of clinical analysis (51).

To examine the contribution of formalin fixation to the outcomes of DNA methylation analysis, we generated FFPEs from flash frozen surgical glioma specimens cryostored in CryoTrays. FFPE blocks were sampled with a microtome, and, after deparaffinization/rehydration, DNA was prepared using PIXUL and analyzed by 5mC MeDIP (29). The 5mC signal distribution from the FFPE samples at the -57bp promoter and enhancer 1 sites largely recapitulated distribution of mean, median and heterolon values generated from the frozen tissues **(Fig.S2C-F).**

Using 5mC MeDIP-seq we have previously shown high genome wide DNA methylation correlation between paired frozen and FFPE mouse organ tissues (R^2^ 0.87-0.92) (29). **Fig.3** illustrates comparison of DNA methylation measured in paired frozen and FFPE surgical samples using MeDIP and MS-PCR at the *MGMT* promoter sites. Violin plots show that DNA methylation levels measured by MeDIP-qPCR across all the paired frozen-FFPE specimens are not statistically different between each other **(Fig.3C).** Side-by-side MeDIP-qPCR comparison using our three-tier stratification criteria shows that about 25% of the frozen-FFPE tissue pairs did not agree (**Fig.3D**), while according to MS-PCR based four-tier stratification (UW clinical lab) about 30% of the frozen-FFPE pairs of samples were discordant (**Fig.3E**). Side-by-side comparison frozen tissues assayed by MeDIP vs. MS-PCR showed that 25% of fragments did not agree (**Fig.3F**), while MeDIP vs. MS-PCR comparison of FFPEs revealed that more than 30% samples differed (**Fig.3G**). The degree of the differences between these two methods is similar to what has been reported for side-by-side comparison of MS-PCR with pyrosequencing (34). The differences in measured *MGMT* promoter methylation levels between frozen and matched FFPE samples might reflect not only technical noise and intratumor heterogeneity but also the differences in metrics used to stratify MeDIP results compared to the MS-PCR criteria used in clinical labs. It remains to be determined if these *MGMT* promoter methylation differences between frozen vs. FFPE and/or MeDIP vs. MS-PCR are clinically relevant. To aid data interpretation, digitized images of H&E stained slides for neuropathology interpretation were prepared from 2-3 1x2mm cores extracted (CryoCore) from frozen tissues and then fixed (FFPE). There were cases where either frozen, FFPE, or both were stratified as *MGMT* promoter methylated but histology was interpreted as negative for tumor. (**Fig.3D-G**). This seeming discrepancy may reflect histological heterogeneity of surgical specimens (see below).

**Fig. 3.**
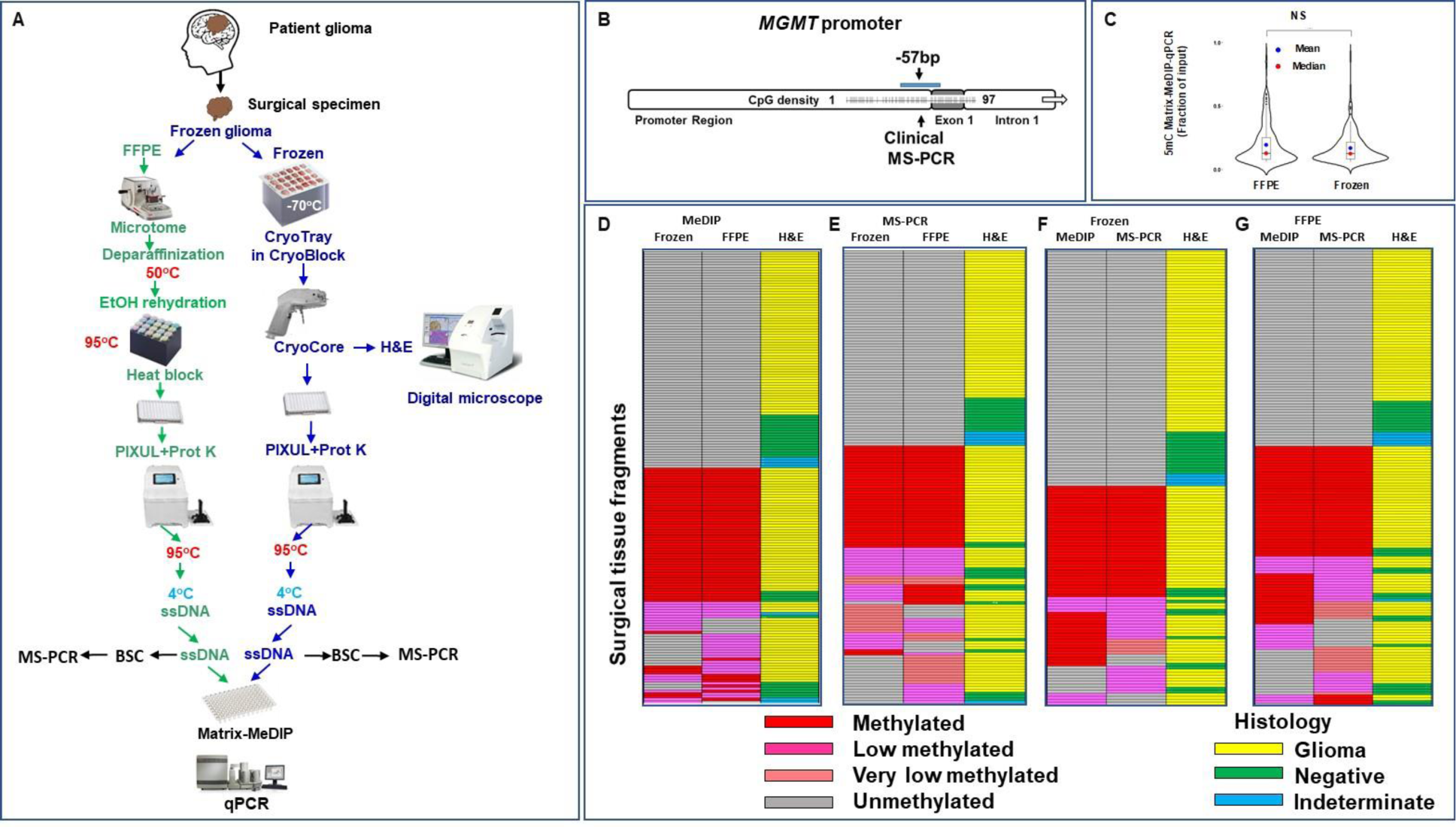
Comparison of *MGMT* promoter methylation MeDIP and MS-PCR data (Methylated, Low methylated, Very low methylated (only for MS-PCR), and Unmethylated) and H&E histology of paired FFPE and Frozen surgically resected sample collection. ***A.*** 96-well plate PIXUL-based protocol was used to isolate DNA from 173 paired FFPE and frozen GBM tissue samples. Matrix-MeDIP was done using 5mC antibody. CryoCore was also used to sample each frozen tissue for H&E digital microscopy and histopathologic interpretation. ***B.*** MGMT promoter CpG sites assayed by qPCR. ***C.*** Violin plots comparing Matrix-MeDIP-qPCR results from matched frozen and FFPE blocks from surgical specimens. ***D-G.*** Surgical specimen heatmaps showing comparisons of stratified *MGMT* promoter methylation status measured by MeDIP and MS-PCR in Frozen and FFPE samples ***(D-E)***. Comparison MeDIP vs. MS-PCR are shown for Frozen ***(F)*** and FFPE ***(G)*** respectively. Two-three ∼1x2mm cores sampled from each tumor fragment were used to prepare H&E slides for digital imaging and histopathological interpretation (Glioma, Negative-no tumor, Indeterminate) matched to *MGMT* promoter status ***(D-G, third column)***.

### Effects of short (24hrs) and long (72 hrs) formalin crosslinking time on GBM DNA (5mC) methylation measurements (**Fig.4**)

Duration of formalin fixation might influence DNA (52,53), but it is not known if and how these conditions affect GBM deamination of 5mC at the *MGMT* promoter. Resected tissues were flash frozen, each specimen was divided into three sections: one was immobilized in CryoTray, while the other two were formalin cross-linked for either 24 or 72 hours and then embedded in paraffin. DNA was isolated from these specimens and used in Matrix-MeDIP with 5mC antibody. The MeDIPed DNA was used in qPCR and to generate sequencing libraries **(Fig.4A)**. **Fig.4B** shows that the 5mC signal distribution at *MGMT* enhancer, promoter, and exon5 (which contains multiple CpGs, **Fig.S1**) sites was not different between the frozen and corresponding formalin cross-linked (24 or 72hr) samples.

**Fig. 4.**
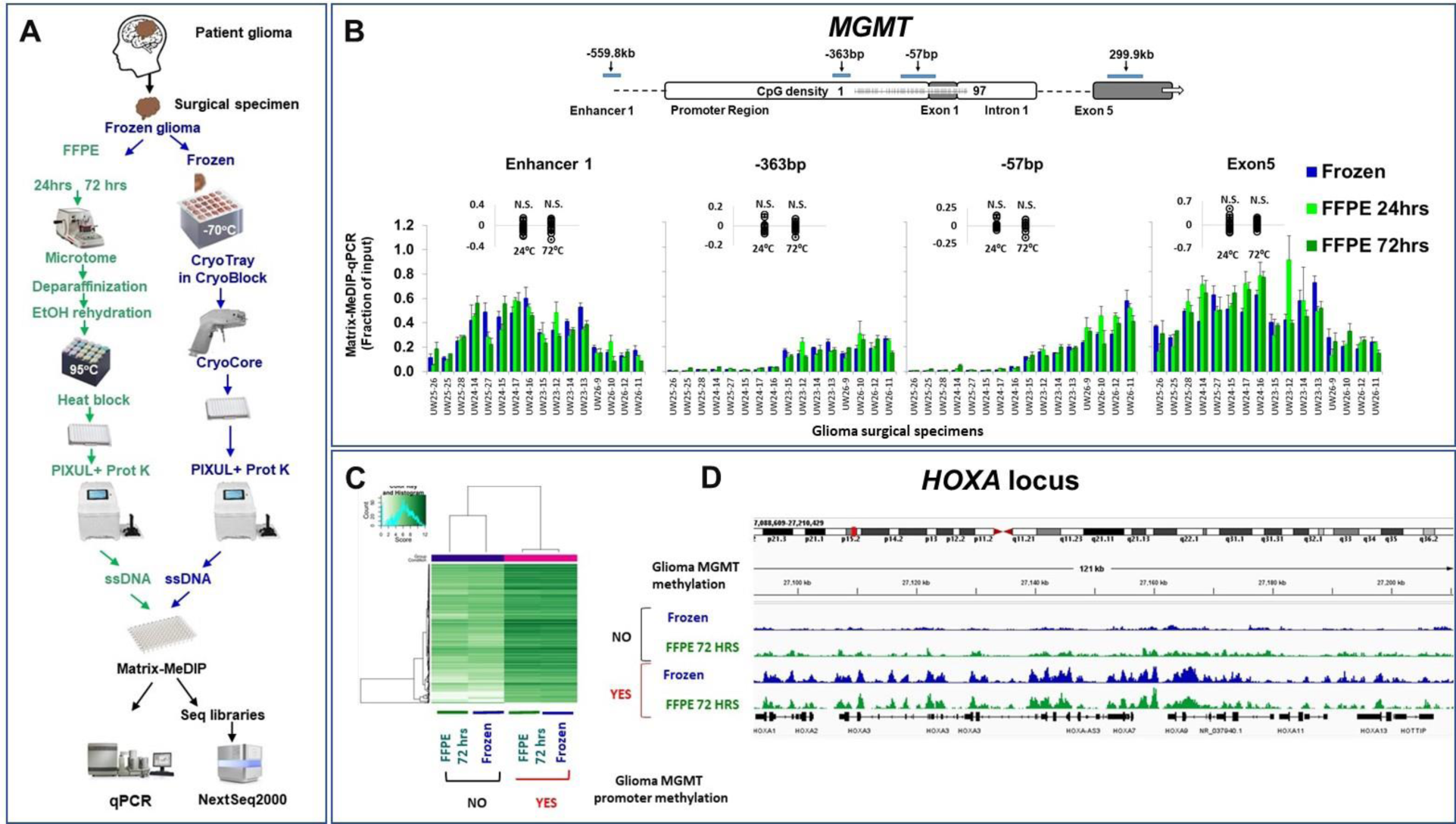
Effects of short (24hrs) and long (72hrs) formalin crosslinking on glioma DNA (5mC) methylation. ***A.*** Resected tumors were flash frozen. Frozen specimens were divided into three pieces: one was immobilized in CryoTray, while the other two were formalin cross-linked for either 24 or 72 hours and embedded in paraffin (FFPE). FFPE samples were sectioned with microtome, and sections were used to isolate DNA using PIXUL. CryoCore was used to sample tissues from the CryoTray, and cores were jetted into 96-well PIXUL plate on ice to isolate DNA. Denatured DNA (ssDNA) was used in Matrix-MeDIP using 5mC antibody (MeDIPed DNA). The MeDIPed DNA was used in qPCR and in the generation of sequencing libraries. Sequencing was done using NextSeq2000. ***B.*** qPCR analysis at the indicated *MGMT* gene sites, Mean ±SEM, three (n=3) different fragments from the same surgical specimen. The inserts above the column graphs show the FFPE (24 and 72 hours) minus Frozen 5mC signal difference for each paired tumor sample (each *circle* represent a difference). ***C.*** The heatmap shows DNA methylation at multiple genomic sites clustering by the methylated (***yes***) and unmethylated (*no*) status of *MGMT* promoter measured in DNA from frozen vs. FFPE specimens. ***D.*** Snapshot was generated from bigwig files using Integrative Genomics Viewer (IGV) (91) along the *HOXA* locus.

To assess epigenome-wide methylation patterns we sequenced 5mC MeDIP DNA from GBM with methylated and unmethylated *MGMT* promoter. This analysis showed that samples cluster by *MGMT* promoter methylation status but not by frozen vs. FFPE **(Fig.4C)**. *HOXA* cluster belongs to a highly conserved *HOX* homeobox class of genes that encode transcription factors required for normal development (54). Aberrant *HOXA* locus expression of sense and antisense *HOX* transcripts (e.g. long non-coding RNA *HOTAIRM1* (55)) has been widely reported as altered in malignancy including DNA hypermethylation in gliomas (40,56). The presented IGV snapshots show that tumors with methylated *MGMT* promoter had multiple methylated regions along the *HOXA* locus that were not found in gliomas with unmethylated *MGMT* promoter gliomas, in both frozen and FFPE specimens. Thus, duration of formaldehyde cross-linking did not appear to affect the levels of the 5mC signals **(Fig.4B-D)**.

### Effects of *ex-vivo* cold and warm ischemia on the methylation status of 5’ *MGMT* locus in GBM-derived mouse xenografts (**Fig.5**)

DNA methylation levels are determined by the balance between activities of DNA methylases (DNMT1, DNMT3a, and DNMT3b) and ten-eleven translocation (TET) family enzymes that oxidize 5-methylcytosine to 5-hydroxymethylcytosine, the first step in active demethylation (57). Expression of DNMT and TET enzymes and their activities are affected by tissue hypoxia (58,59). Importantly, hypoxia can alter DNA methylation *in vivo* (59,60). Further, after the blood supply is cut off, the *ex-vivo* tissue is under hypoxic/nutrient deprivation stress followed by waste accumulation, which can cause profound cellular and molecular changes (61). Cooling of tissues or organs slows down tissue damage, a practice used in organ procurement for transplantation but less so with resected tumor specimens, which are often left at room temperature in the operating room. *Ex-vivo* ischemia can affect gene expression(62) but it is unknown whether or not it alters *MGMT* promoter methylation. Also, 5mC is prone to spontaneous non-enzymatic deamination to generate thymine (63,64) skewing measurements.

**Fig. 5.**
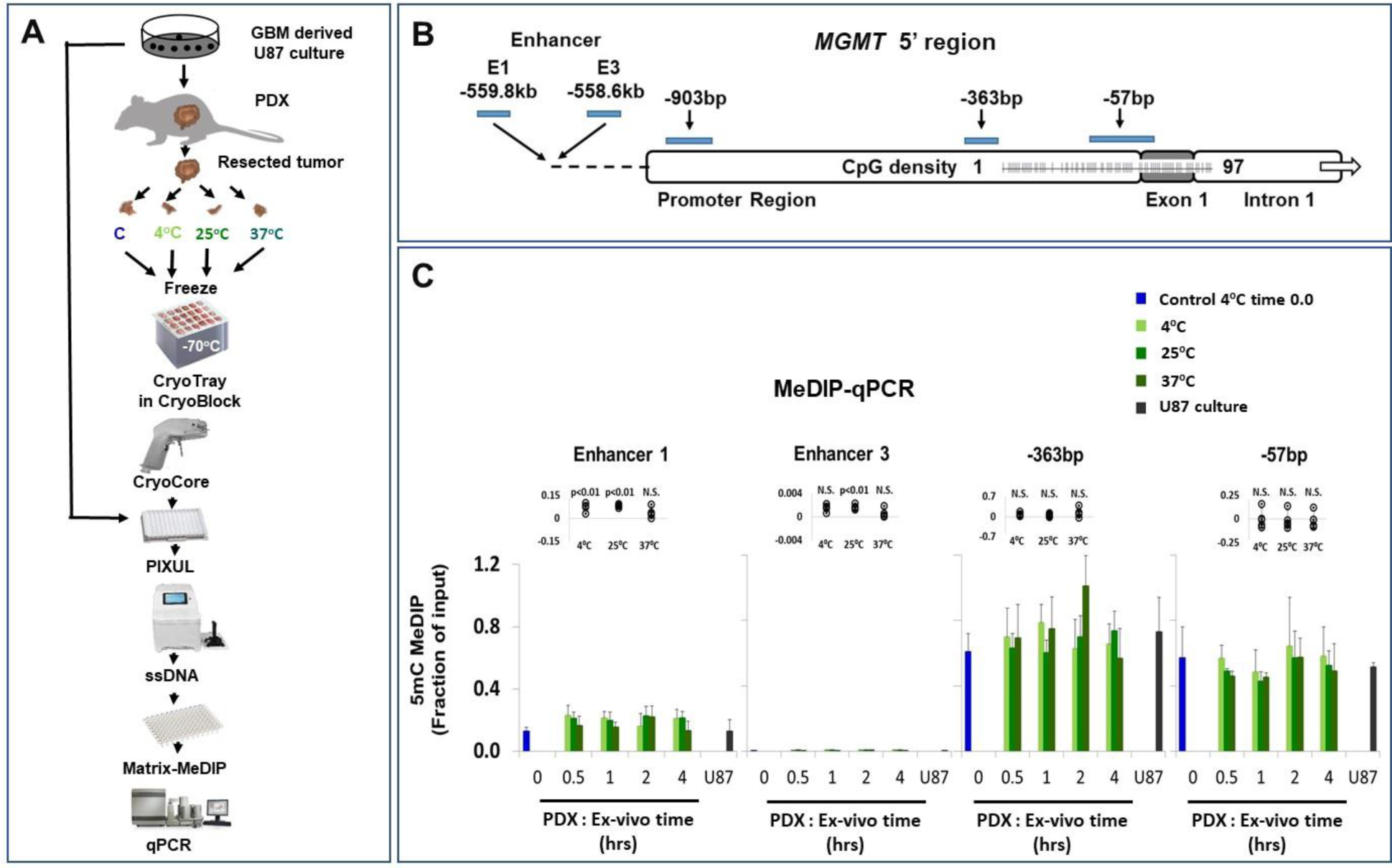
Effects of ex-vivo cold and warm ischemia on methylation of 5’ *MGMT* region in GBM-derived xenograft tissues in mouse. ***A.*** Xenograft tissues were generated by engrafting GBM-derived cultures (U87 (25)) in the flanks of nude mice. After resection, each xenograft tissue was divided into four pieces and processed as follows: one piece was immediately flash frozen and the other three were kept at 4, 25, and 37°C, with each of the three pieces being further divided for incubations of 0.5, 1, 2, or 4hrs at their respective temperatures – a total of 13 different conditions. After the given time points, all samples were flash frozen. All flash frozen samples were then cryostored (-80°C) in CryoTrays. Tissue cores extracted with the CryoCore were ejected into a 96-well plate and treated in PIXUL to isolate DNA. In parallel, DNA was isolated from the parental GBM-derived U87 pellets. ssDNA was used in Matrix-MeDIP-qPCR using 5mC antibody. ***B.*** The 5’ MGMT region sites assayed by PCR. ***C.*** Matrix-MeDIP-qPCR results, Mean±SEM, n=4. The inserts above the column graphs show the difference (*circles*) of experimental minus control (shown as 0 time point) 5mC signal difference for each sample.

Testing the effects of time course of *ex-vivo* ischemia in the operating room on glioma *MGMT* promoter methylation analysis is logistically challenging, limiting the number of samples that can be collected. In addition, intratumor heterogeneity could complicate interpretation of the data. Before doing these studies on surgical samples, we used GBM-derived cell culture xenografts in nude mouse models (**Fig.5A)** to define the effects of *ex-vivo* ischemia on *MGMT* promoter methylation in the resected xenograft tissue. After resection, each xenograft tissue was divided into four pieces and processed as follows: one piece was immediately flash frozen and the other three were kept at 4, 25, and 37°C, with each of the three pieces being further divided for incubations of 0.5, 1, 2, or 4hrs at their respective temperatures – a total of 13 different conditions. After the given time points, all samples were flash frozen. All frozen samples were then cryostored (-80°C) in CryoTrays. Tissue cores extracted with the CryoCore were ejected into 96-well plate wells and treated in PIXUL to isolate DNA. In parallel, DNA was isolated from the parental GBM-derived U87 pellets (25). ssDNA was used in 5mC Matrix-MeDIP-qPCR analysis of the *MGMT* promoter and enhancers regions **(Fig.5B)**. The results demonstrate that warm ischemia at all temperatures including 37°C up to 4hrs had no significant effect on *MGMT* promoter methylation levels **(Fig.5C)**. We used these results to plan *ex-vivo* testing in the operating room.

### Effects of *ex-vivo* warm ischemia on resected brain tumor tissue 5mC methylation levels (**Fig.6**)

In the operating room 4 hours represents a practical upper limit for assessing pre-analytical conditions on *MGMT* promoter methylation. Surgically resected tissues were divided: one fragment was snap-frozen while the other was kept at 37°C for 4hrs in an incubator. Both surgical fragments were then cryostored (-80°C) in CryoTrays. CryoCore was used to sample tissues from the CryoTray. DNA was isolated with PIXUL, and ssDNA was used in Matrix-MeDIP using 5mC antibody. The MeDIPed DNA was used in qPCR and sequencing analyses **(Fig.6)**. The MeDIP-qPCR results show that there was little or no differences in the 5mC levels between frozen and *ex-vivo* warm ischemia samples at the *MGMT* promoter, enhancer 1, and exon5 sites **(Fig.6B).** Genome-wide DNA methylation analysis showed that the sequencing data cluster by *MGMT* promoter methylation status and not by frozen vs *ex-vivo* warm ischemia conditions **(Fig. 6C).** IGV snapshot shows that there is hypermethylation along the *HOXA* locus in *MGMT* promoter-methylated samples that is also not affected by *ex-vivo* ischemia **(Fig. 6D).**

**Fig. 6.**
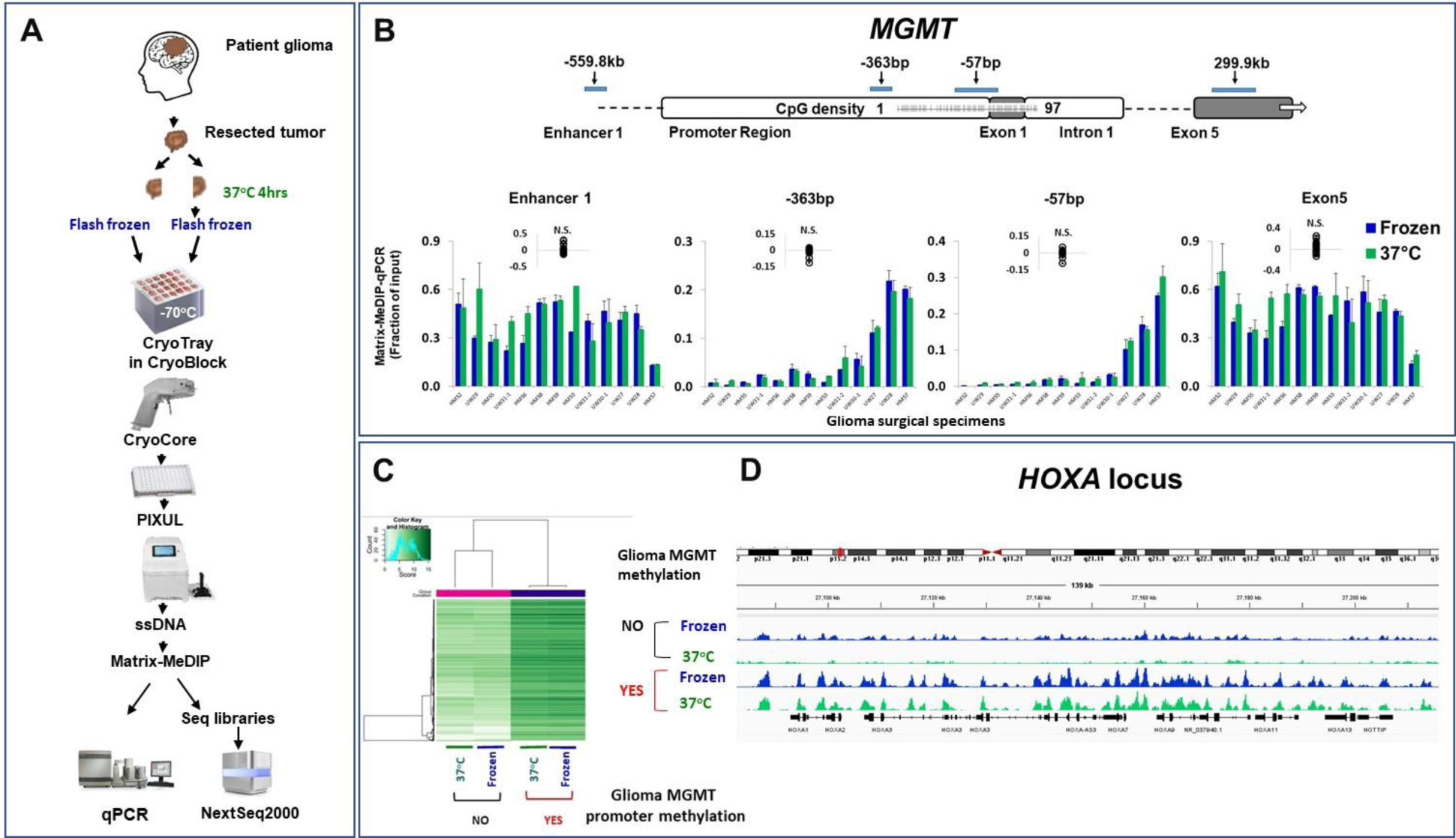
Effects of *ex-vivo* warm ischemia on glioma DNA (5mC) methylation. ***A.*** Resected tumor was divided into two fragments: one was snap frozen while the other was kept at 37°C for 4hrs in an incubator. Both surgical fragments were then cryostored (-80°C) in CryoTray. CryoCore was used to sample tissues from the CryoTray, and cores were jetted into 96-well PIXUL plate on ice. DNA was isolated with PIXUL, and ssDNA was used in Matrix-MeDIP using 5mC antibody. The MeDIPed DNA was used in qPCR and in the generation of sequencing libraries. Sequencing was done using NextSeq2000. ***B.*** qPCR analysis at the indicated MGMT gene sites, Mean ±SEM, three (n=3) different fragments from the same GBM surgical specimen. The inserts above thc column graphs show 5mC signal difference between the 37°C and the frozen state (*circles*) for each paired tumor sample. ***C.*** The heatmap shows that the MGMT promoter methylated (***yes***) and unmethylated (*no*) DNA methylation sites cluster together much more closely than the data from the 37°C vs. frozen samples. ***D.*** Snapshot was generated from bigwig files using IGV, along with the *HOXA* locus.

### Effects of repeated freezing and thawing on resected brain tumor DNA methylation (**Fig.7**)

Cryopreservation is a common way to store and transport tissue samples. Freezing and thawing could also affect DNA integrity and methylation status. Although it is not uncommon to thaw frozen tissue samples for analysis, the effect of repeated cycles of freezing and thawing of gliomas on DNA methylation is not known. Fragments of frozen glioma specimens were subjected to three cycles of freezing-thawing, and, after the last freeze, were sampled along with the original specimens that were kept frozen. DNA was isolated and assayed by Matrix-MeDIP-qPCR/seq **(Fig.7A)**. qPCR results demonstrate that repeated cycles of thawing and freezing of surgical specimens do not alter the 5mC levels at the *MGMT* promoter, enhancer 1, and exon 5 sites **(Fig.7B).** Genome-wide DNA methylation analysis shows that the sequencing data clusters by *MGMT* promoter methylation status but not by freezing-thawing **(Fig.7C).** Previous studies indicated that mitochondrial dysfunction may contribute to altered glioma energy metabolism (65) and may contribute to TMZ resistance (66). Mitochondrial dysfunction has also been linked to altered mitochondrial DNA (mtDNA) copy number and methylation status in gliomas (65). Given that mtDNA is methylated, mitochondrial epigenetic processes during thawing could alter mtDNA methylation. Representative IGV snapshots show that in this set of samples there was a higher level of mtDNA methylation in *MGMT* promoter unmethylated glioma samples compared to methylated, but that repeated freezing and thawing did not alter mtDNA methylation **(Fig.7D).**

**Fig. 7.**
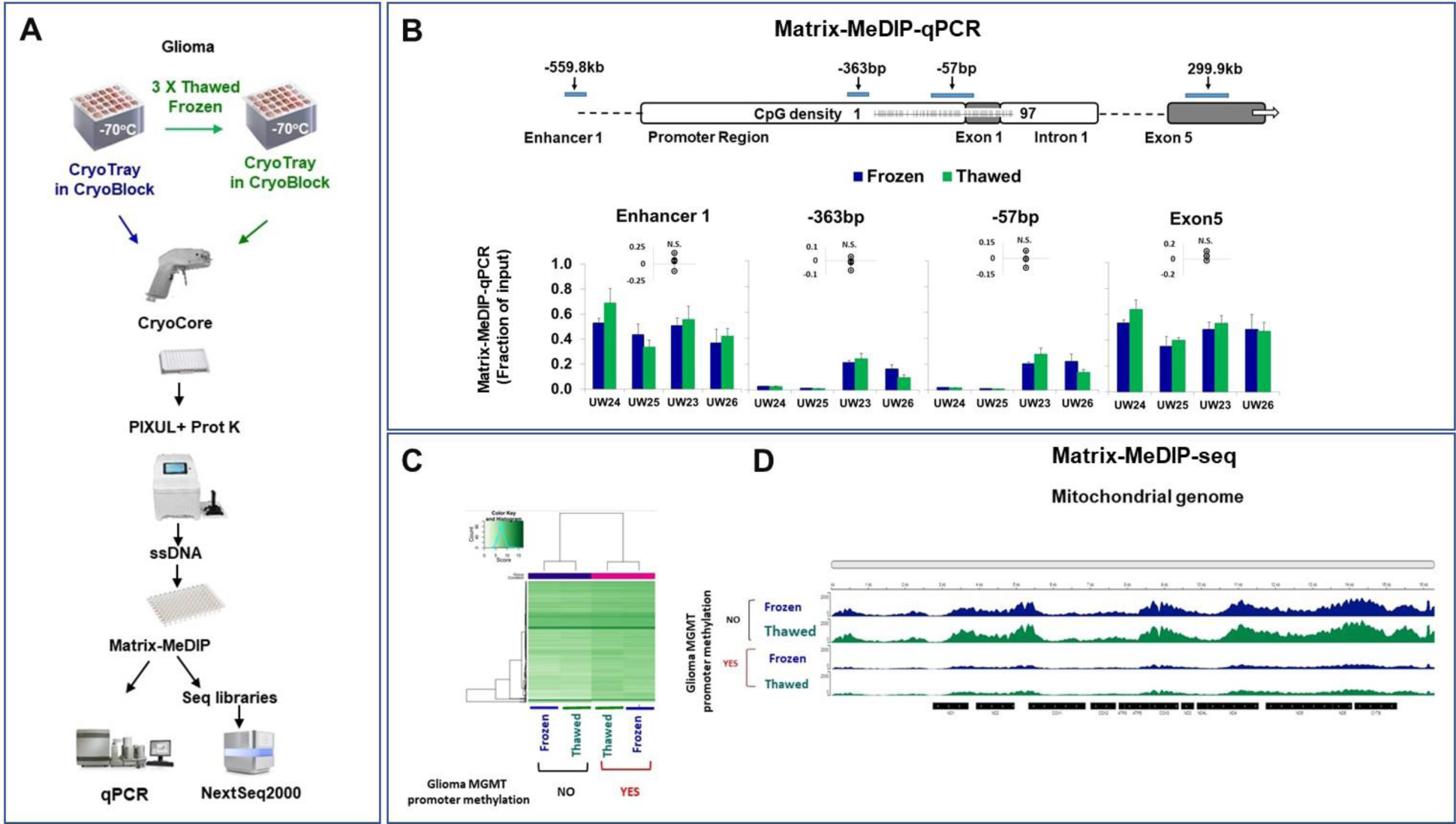
Effects of repeated freezing and thawing on glioma DNA (5mC) methylation. ***A.*** Three different frozen fragments in CryoTray from the same surgical glioma biospecimen were sampled with the CryoCore, and cores were jetted into 96-well PIXUL plate on ice. These tissue fragments were then thawed and placed in a new CryoTray and freeze-thawed three times. After the last freeze, the frozen tumor fragments were sampled with the CryoCore and cores were jetted to the 96-well PIXUL on ice. DNA was isolated with PIXUL, and ssDNA was used in Matrix-MeDIP using 5mC antibody. The MeDIPed DNA was used in qPCR and in the generation of sequencing libraries. Sequencing was done using NextSeq2000. ***B.*** qPCR analysis at the indicated *MGMT* gene sites, Mean±SEM, three (n=3) different fragments from the same surgical specimen. The inserts above the column graphs show the Thawed minus Frozen 5mC signal difference (*circles*) for each tumor sample. ***C.*** The heatmap shows that the *MGMT* promoter methylated (***yes***) and unmethylated (*no*) DNA methylation sites cluster together more closely than the data from the three times frozen-thawed samples. ***D.*** Snapshot was generated from bigwig files using IGV along the mitochondrial genome.

The above studies suggest that DNA methylation is a robust chemical modification in gliomas that is in agreement with other prior studies. Kopfnagel et.al. showed that after 10 DNA freeze and thaw cycles no significant difference in methylation of examined CpGs sites (67). Lee et.al., found that after cryopreservation of DNA for several years there is no detectable change in mean global DNA methylation, there was detectable bias towards hypomethylation in 4,049 CpGs out of 830,545 CpGs tested, or in only 0.5% of the probes (68). Li et.al., provided evidence that methylation profiles of defined genomic regions in the DNA or whole blood samples archived for 20 years were similar across all storage temperatures, including storing at 4°C(69). Sasaki et.al., reported that maintaining whole blood *ex vivo* for up 2 hrs at 4°C or dried blood spots up to a week at room temperature had little effect on DNA methylation profiles (70).

### Heterogeneity of glioma *MGMT* promoter methylation (**Fig.8**)

Increased understanding of the molecular mechanisms that underlie tumor heterogeneity shifts diagnostic and therapeutic paradigms in that multiple, rather than single, biopsies may be needed to optimize personalized care (71–74). DNA methylation exhibits spatially heterogeneous patterns within a given GBM (9,12,30,75–77). This raises questions about the degree of intratumor heterogeneity of *MGMT* promoter methylation and its correlation to histological landmarks and *IDH1/2 (isocitrate dehydrogenase)* mutations (6,78) that affect GBM patient survival and response to therapy. The presence or absence of intratumor heterogeneity of *MGMT* promoter methylation could reflect tumor/biopsy size, and perhaps most importantly match histological heterogeneity (79). Thus, intratumor heterogeneity is a critical pre-analytical factor that warrants in-depth investigation. Yet the standard practice is to assay *MGMT* promoter methylation from a single biopsy, regardless of the tumor size and histology. Given the frequently observed geographic intratumor heterogeneity, this practice could be misleading for patient stratification and management.

**Fig. 8.**
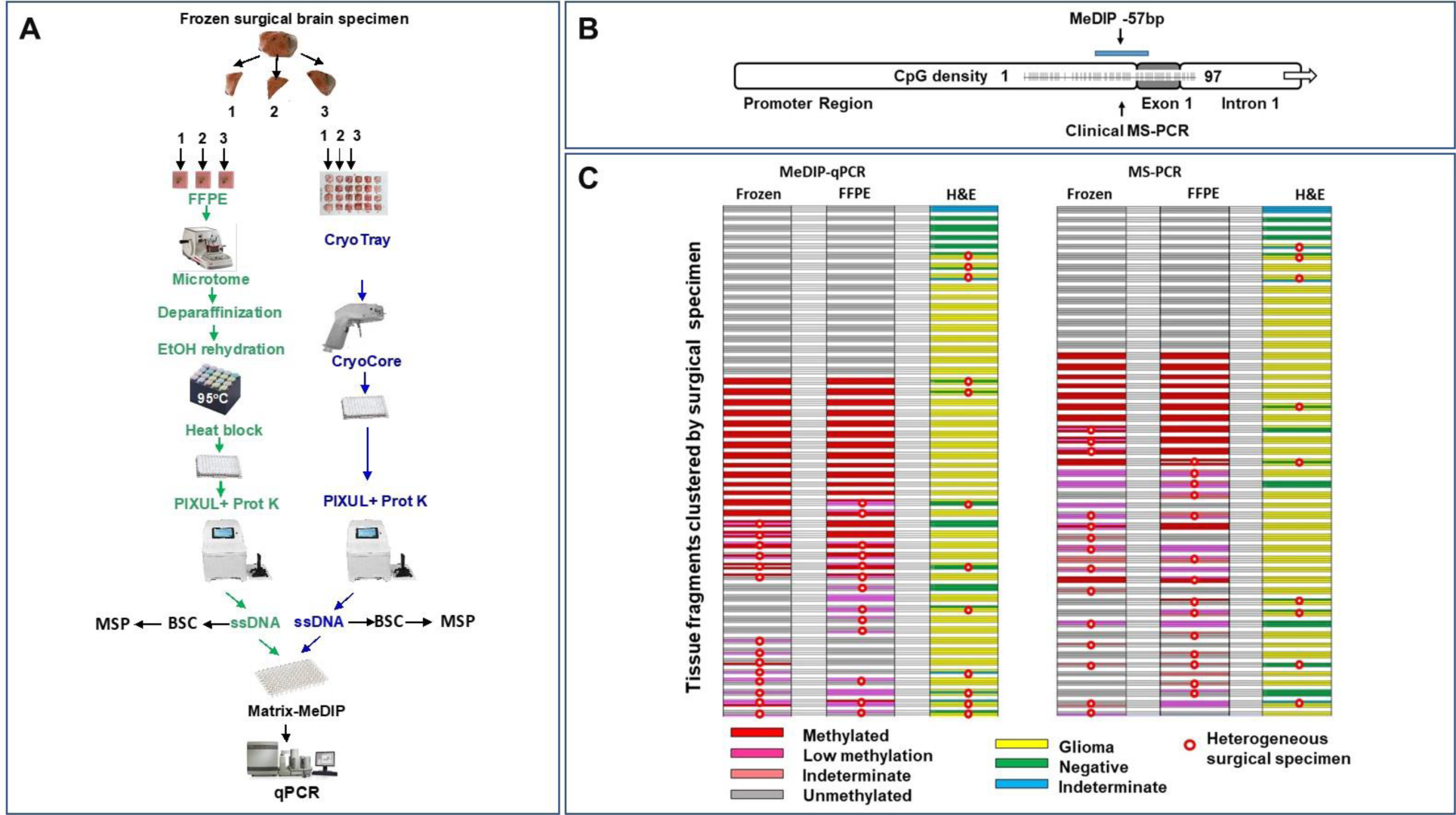
Heterogeneity of GBM *MGMT* promoter methylation by stratified metrics. ***A.*** Surgical specimens were flash frozen and then divided into 2-3 sections while still frozen. Each frozen section was divided into approximately equal pieces: one was used to make an FFPE block while the other was placed in a CryoTray maintained at less than -70°C in CryoBlock. ssDNA was generated from frozen and FFPE tissues, and DNA metylation was assessed by Matrix-MeDIP-qPCR and bisulfite conversion (BSC)-MS-PCR. Two-three ∼1x2mm cores sampled from each fragment were used to prepare H&E slides for digital imaging and histopathological interpretation (Glioma, Negative-no tumor, Indeterminate) ***B***. Location of the PCR primers. ***C.*** Heatmaps of stratified *MGMT* promoter methylation status tissue fragments clustered by the same surgical specimen. Rows of colored fields representing 2-3 fragments derived from the same surgical specimen are clustered together and are separated by white field from the grouped fragments derived from other tumors. The red circles designate surgical samples where one tissue fragment yielded a different result from the other fragment(s), i.e., heterogeneity.

Using the CryoGrid-CryoCore system integrated with the PIXUL sample preparation platform allows one to analyze multiple geographic sites of different tissue fragments generated from the same tumor. This process generating dozens of core biopsies is faster and their analysis is less costly compared to other the currently available methods. Surgical specimens were flash frozen and then divided into 2-3 sections while still frozen. Then, each frozen section was divided into two approximately equal pieces: one was used to make an FFPE block while the other was placed in CryoTray (-80°C) in CryoBlock. DNA was isolated as above using PIXUL and assayed by both MeDIP and MS-PCR **(Fig.8A).** We collected 50 surgical specimens that were large enough to divide into two or more sections. Using the above criteria for unmethylated, intermediate, and methylated *MGMT* promoter assayed by MeDIP-qPCR, there were 30% and 26% heterogenous (i.e. at least one section different from the other section(s) of the same surgical specimen) frozen and matched FFPE tissues, respectively. With the same DNA assayed by MS-PCR using the criteria unmethylated, indetermined, low methylated, and methylated, there were 34% and 32% heterogenous frozen and FFPE tissues, respectively. Intratumor *MGMT* promoter methylation heterogeneity is a well described observation in gliomas, but the frequency in any given study varies. Verburg et.al. found that *MGMT* promoter methylation, based on two probes per tumor, was found in 30% of patients and was not related to tumor purity, *IDH1/2* mutation status, or histology (12). Gempt et.al. recently reported that 16% of their tumor samples contained both methylated and unmethylated *MGMT* promoter (74). Yet Wegner et.al. found that only 1/11 patients with *IDH* wt and 1/1 *IDH* mutated tumors were heterogeneous for *MGMT* promoter methylation (75). Differences in methods used may account for some but not all of the discrepancies given that some studies used the same EPIC platform (12,74,75). The reasons underpinning this glioma intratumor *MGMT* promoter methylation heterogeneity thus far is unclear.

Intratumor histological heterogeneity is a hallmark of gliomas (79,80). Previous studies have underscored that histological tumor features could be under-represented when the size of tested tissues on H&E slides is small (79). Also, the resected tumor may contain variable amounts of normal tissues which may further confound histological interpretation. Based on H&E whole slide digital images assessed as negative, undetermined, and glioma, 24% of surgical specimens were histologically heterogeneous **(Fig.8C).** The above studies suggest that our CryoGrid system, which allows multiple sampling, high throughput, and low cost of PIXUL-Matrix-MeDIP assays, offers an alternative way to assess intratumor *MGMT* promoter and genomewide DNA methylation heterogeneity along with histopathology (23,29).

### Differential DNA methylation analysis uncovers association of *MGMT* promoter methylation status with DNA methylation and transcriptional repression of long non-protein coding RNA (lncRNA) (Figs.9 and S3)

We found that MGMT promoter methylation status clusters with DNA methylation of other loci (**Figs.4C, 6C,** and **7C**). Differential DNA methylation of samples stratified as *MGMT* promoter methylated and unmethylated and Gene Ontology (GO) analysis uncovered association with methylation of lncRNA loci. Altered expression of lncRNAs has been implicated in tumorigenesis including gliomas (81–83). **Fig.S4** shows IGV snapshot of DNA methylation profiles of several lncRNA loci that correlate positively (*LINC02106, LINC01096, LINC02875 LINC01150, LINC01305, HOTAIR lncRNA*, *MIR1915HG*) and negatively (*LINC01526*) with *MGMT* promoter methylation status. There is evidence that expression of some of these could serve as glioma prognostic biomarkers, e.g., *HOTAIR* (83)*, LINC02875* and *LINC01305* (81). Notably, serum miR-1915-3p, encoded by *MIR1915HG* host gene, has shown biomarker utility in classifying patients with diffuse glioma (84). Altered expression of *LINC02106* has not been previously reported. **Fig.9** shows that, in the collection of gliomas that we tested, inverse relationship of DNA methylation and mRNA expression of *LINC02106* and *MIR1915HG* mirror *MGMT* promoter methylation and gene expression **(Fig.9B-G).** Given lncRNA involvement in multiple tiers of gene expression (85–87), their emerging prognostic value (83) and the pre-analytical resilience, DNA methylation of lncRNA genes offers exciting avenues to advance biomarker research in GBM.

**Fig. 9.**
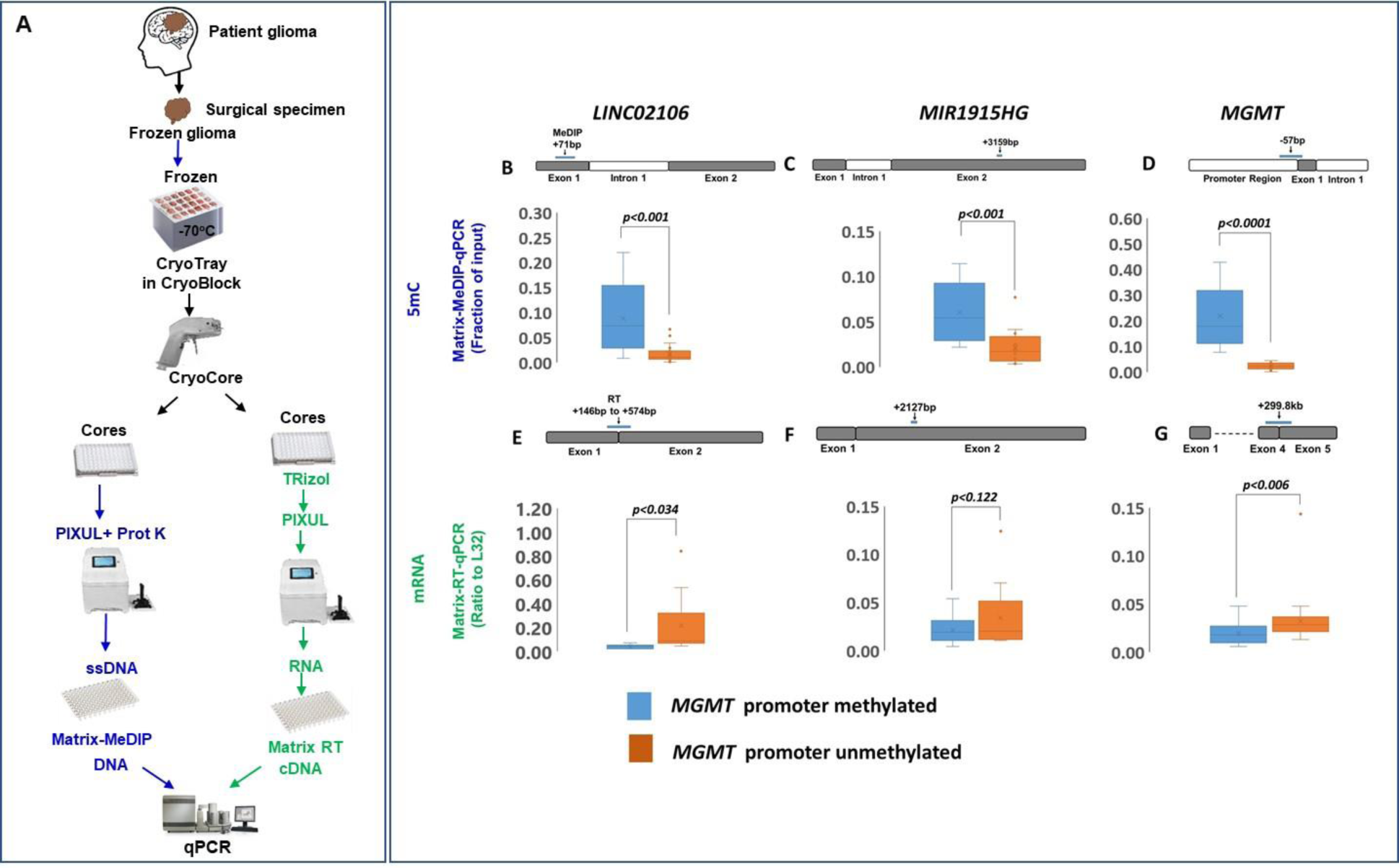
DNA methylation and mRNA expression of long non-protein coding intragenic RNA (lincRNA), *LINC02106 and MIR1915HG,* and *MGMT* genes in gliomas with methylated (*blue*) and unmethylated (*orange*) *MGMT* promoter. ***A.*** Frozen gliomas in CryoTray were sampled with the CryoCore, with cores jetted into two 96-well plates: one to isolate DNA and the other to isolate RNA using PIXUL. ssDNA was used in 5mC Matrix-MeDIP, and RNA in Matrix-RT to generate cDNA. DNAs were used in qPCR using primers to *LINC02106, MIR1915HG* and *MGMT* as shown for 5mC Matrix-MeDIP (Fraction of input) (***B*** and ***D***) and mRNA Matrix-RT (Ratio to *L32*) (***E*** and ***G***). Gene cartoons, white designates promoter or intron regions, black exons.

In summary, this study demonstrates the following, i) 5mC *MGMT* promoter methylation levels estimated by MeDIP-qPCR and MS-PCR methods highly overlap suggesting that MeDIP can be used as an alternative approach. Moreover, MeDIP combined with sequencing allows to assess 5mC DNA methylation genome wide which MS-PCR cannot. ii) In agreement with other studies (67–69), 5mC DNA methylation status exhibits resilience to sample handling variations that may occur in the operating room during collection of surgical specimens, their storage and laboratory processing. As such, DNA methylation is a robust biomarker as compared to some others that are more labile (e.g. phosphoprotein (88)). iii) FFPE tissues, which are convenient to store and transport, can be used reliably to assess DNA methylation at specific loci and/or genome wide (29). 5mC modification is also chemically quite durable (29,89), given it survives the harsh fixation used to prepare FFPEs, long storage at room temperature, and high temperature treatments during DNA retrieval. iv) GBM exhibits *MGMT* promoter methylation intratumor heterogeneity suggesting that multiple geographic sites of resected tissues need to be biopsied for patient stratification. In this regard, the CryoGrid-PIXUL-Matrix-MeDIP (23,29) is a well-suited platform to analyze multiple regions of frozen or FFPE surgical tissues. v) *MGMT* promoter methylation status is associated with differentially methylated loci including *HOXA*, *lncRNA* and miRNA genes, opening new avenues to explore more granular data-based biomarkers to further advance glioma patients’ stratification.

There are other questions that warrant further investigation. i) What is the clinical significance of *MGMT* promoter methylation in GBM intratumor heterogeneity? For example, what is the clinical impact when there is only one out of several, geographic GBM location that is methylated? ii) What is the optimal number of geographic sites that should be biopsied and how does this number depend on tumor size? iii) Could there be DNA methylation markers (e.g. *lncRNA, HOXA*) that can be used to stratify GBM patients that exhibit little or no intratumor heterogeneity? In this regard, the current and other studies (40) identified many other differentially methylated loci that could be considered to test as biomarkers in individuals diagnosed with gliomas (83,90).

## DECLARATIONS

### Ethics approval and consent to participate

#### Authors’ Contributions

KB and RCR conceived the project, designed the experiments, analyzed the data, and wrote the paper. DM designed, carried out experiments, analyzed the data, and wrote the paper. OD assisted in experimental design and edited the paper. SP interpreted digital H&E histopathology. MV carried out sample collection, processing, generating cell lines and performing xenograft experiments and edited the paper. JD, AP, and RGE provided tissues and edited the paper. RR assisted in experimental designed and edited the paper.

#### Inclusion and Diversity

One or more of the authors of this paper self-identifies as an underrepresented ethnic minority in science. One or more of the authors of this paper self-identifies as a member of the LGBTQ+ community. One or more of the authors of this paper self-identifies as living with a disability. One or more of the authors of this paper received support from a program designed to increase minority representation in science.

## ACKNOWLEDGMENTS

This work is dedicated to Susan B. who passed away from glioblastoma, a dear family friend (KB). We thank UW Laboratory Medicine and Pathology Genetics and Solid Tumors Laboratory for running the MS-PCR assays.

## SUPPLEMENT

PIXUL-Matrix-MeDIP based analysis of gliomas DNA methylation: Assessment of pre-analytical variables.

## MATERIALS, DEVICES, BIOSPECIMENS, AND METHODS

### MATERIALS

Kits/enzymes (**Table S1**) catalog numbers and commercial suppliers are listed in the supplementary tables.

### Reagents

Accutase Cell Dissociation Reagent (StemPro A1110501), Accumax solution (StemPro A7089), B-27 Supplement (17504044), EGF (PeproTech AF-100-15), FGF (PeproTech 100-18B), N-2 Supplement (17502048), and Neurobasal Medium (Gibco 21103049) were from Thermo Fisher Scientific. cOmplete Mini tablets (Roche 04693159001), Dithiothreitol (DTT) (D0632), EDTA (E3134), Tris–HCl (T3253), IGEPAL CA-630 (I8896), sodium deoxycholate (30970), and tetramethylammonium bromide (TMAB) from Sigma-Aldrich. Acetonitrile (J.T. Baker 9017-03), Bluing Reagent (Epredia 7301), chloroform (J.T. Baker 9180-01), Clarifier 1, 2 (Epredia 7401), Dulbecco’s Modified Eagle Medium (DMEM) (Hyclone SH30021.0), DMEMF12 (Hyclone SH30023.01), Eosin-Y with Phloxine (Epredia 71304), FBS (Hyclone SH30910.03), hematoxylin (Epredia 72711), isopropanol (Acros Organics 3223-0010), phosphate buffered saline (PBS) 10X (Gibco 70013-032), sodium chloride (NaCl S-271-3), Trifluoroacetic acid (TFA) (A116-50), Triton X-100 (BP151), and xylene (X3P1GAL) from Fisher Scientific. NP40 (198596) from MP Biomedicals. Penicillin/streptomycin (P/S 15749) from Invitrogen, fetal bovine serum (FBS 43635-500) from Jr. Scientific, and phosphate buffered saline (PBS 70013-032) from Invitrogen. Ethanol (2716) from Decon Labs.

### Buffers

Preparations of all buffers and stock solutions were done with nuclease free reagents and ultrapure distilled RNAse/DNAse free H_2_O. PBS: 137 mM NaCl, 10 mM Sodium phosphate, 2.7 mM KCl, pH 7.4; TE: 10 mM Tris-HCl, 1 mM EDTA, pH 7.5; Immunoprecipitation (IP) buffer: 150 mM NaCl, 50 mM Tris–HCl (pH 7.5), 5 mM EDTA, NP-40 (0.5% vol/vol), Triton X-100 (1.0% vol/vol); Elution buffer-Proteinase K: 25mM Tris Base, 1% IP Buffer, 1mM EDTA, 80μg/ml Proteinase K; Proteinase K buffer (PK buffer): 10mM Tris-HCl pH 8.0, 10mM EDTA, 0.5% SDS, 500μg/ml Proteinase K, and 40mM DTT.

### DEVICES

#### CryoGrid system

To store and sample large numbers of biospecimens for multi-omics analysis, we designed a platform for cryostoring multiple tissue samples and engineered a hand-held rotary tool for rapid sampling of frozen tissues, CryoGrid system, which consists of CryoBox, CryoBlock, thermometer/thermocouple, QR barcoded CryoTrays, and CryoCore (23).

### PIXUL

A 96-well plate sample preparation sonicator for multi-omics applications (Matchstick Technologies, Inc, Kirkland, WA and Active Motif, Carlsbad, CA) (24).

### BIOSPECIMENS

#### Glioma specimen collection

Glioma patients were included according to protocols approved by the Institutional Ethical Review Boards at the Houston Methodist Research Institute (HMRI, Houston, TX) and at the University of Washington. All patient tissue and blood specimens were collected after receiving informed written consent from patients and according to the principles of Declaration of Helsinki. Tissue specimens were collected immediately after surgery, flash frozen stored at -80°C freezer, and shipped frozen in dry ice for analysis.

### GBM-derived cultures

Two to five mg of GBM tissue specimen were homogenized in Accutase™ Cell Dissociation Reagent for single cell isolation. Cells were resuspended in Accumax™ solution and strained using 40 µM filter. After washing twice with 1X PBS, cells were cultured on laminin coated plate in Neurobasal Medium. All cells were maintained under aseptic conditions in serum-free Neurobasal Medium supplemented with N2, B27, 40ng/µL FGF, and EGF or in DMEMF12 and 10% FBS.

### Xenograft models

All animal studies were performed according to protocols approved by the Institutional Animal Care and Use Committee (IACUC) at HMRI. Four weeks old female NU/J mice (Homozygous for Foxn1) were purchased from Jackson laboratory (Jackson Labs Cat. No. 002019). U87-MG cell lines (2 million cells per animal) were injected subcutaneously at the flank of nude mice (n=3). At end point, mice were euthanized, and tumor tissues were harvested. At time 0 min, a piece of tissue was frozen down for baseline measurement. Ex-vivo cold, ambient, and warm ischemia conditions were maintained by incubating tissue specimen at 4°C, room temperature (RT), and 37°C respectively in PBS buffer. Tissue specimens were collected at 0.5hr, 1hr, 2hr, and 4hr for each condition and then flash frozen and shipped on dry ice for analysis. All samples were shipped to UW by maintaining frozen conditions for downstream analysis.

### METHODS

#### Frozen tissue storing and sampling

To store and sample tissues, we used a platform for cryostoring multiple tissue samples and engineered a hand-held rotary tool for rapid sampling of frozen tissues, CryoGrid system, which consists of CryoBox, CryoBlock, thermometer/thermocouple, QR barcoded CryoTrays, and CryoCore. Frozen tissues are placed in the wells and immobilized with either OCT or Leica CryoGel. The CryoTray, with tissues covered by a QR code labeled lid, is stored at -80°C (23). Tissues are sampled with CryoCore and cores are jetted into tubes or 96-well PIXUL plates.

#### Tissue formalin fixing and paraffin embedding (FFPE), microtome sectioning, and DNA extraction from rehydrated samples

Frozen tissues were fixed with 10% formalin overnight and then embedded in paraffin. Following fixation, the tissue samples were processed through dehydration, clearing, and infiltration. Dehydration was initiated in 70% ethanol, progressing through two changes of 95% ethanol and then three changes of 100% ethanol. The tissues were then cleared by three changes of xylene and infiltrated by melted paraffin wax. FFPEs were sectioned using a microtome, and slices were deparaffinized and rehydrated using serial ethanol dilutions.

#### Hematoxylin and Eosin (H&E) slide preparation and digital image analysis

Paraffin embedded blocks were sectioned on a Leica RM2255 at 4-5 µm, air dried at room temperature overnight, and baked at 60°C prior to staining. Slides with paraffin sections were processed using the Sakura Tissue-Tek Prisma automated stainer for Hematoxylin and Eosin (H&E) staining. Slides were deparaffinized and rehydrated in multiple changes of each of the following reagents: xylene for 5 minutes, 100% ethanol for 4 minutes, 95% ethanol for 4 minutes, water rinse for 1 minute. Slides were stained in hematoxylin for 11 minutes followed by a 1 minute water rinse, a 1 minute clarifier incubation, a 2 minute water rinse, Bluing Reagent for 1 minute, a 1 minute water rinse, 95% ethanol for 1 minute followed by Eosin Y Phloxine for 2 minutes. Slides were dehydrated in graded ethanol with two 1 minute 95% ethanol incubation and 3 100% ethanol incubations and cleared with xylene for at least 4 minutes. Whole slide digital imaging was performed on an Aperio AT Turbo or Aperio Versa 200 at 20X. Once reviewed, an Excel spreadsheet was generated of the diagnoses of each individual slide and the file was submitted for archiving and correlation.

#### DNA isolation from frozen tissues

One CryoCore core extracted from frozen tissue is jetted with PBS (drawn from the CryoCore syringe reservoir) directly into 1.5ml tubes on ice. PBS is removed and 100μL of 25mM Tris Base pH 10.2 + 1uL IP buffer + 3uL RNase (10mg/mL stock) is added to the tubes, which are incubated for 30min at RT. 4μL Proteinase K (20mg/mL stock) is added to each tube, and contents are transferred to wells of 96-well plate and processed in PIXUL (no pre-chilling, Pulse=50, PRF=1.0, Burst=20, T=30min). Plates are briefly centrifuged, and content is transferred to 1.5mL tubes which are then boiled for 10min, chilled on ice, and centrifuged at 16,000xG for 15min. The supernatants containing ssDNA are used immediately in MeDIP or can be stored frozen (-20°C) for future use.

#### DNA isolation from FFPE tissue blocks

10-20 thin slices (5μm) are cut with microtome, collected in 1.5mL tubes. Paraffin is removed using SafeClear (1.2mL) at RT (10min) and tubes are centrifuged at 10,000xG for 2min. Samples are rehydrated with serial dilutions of 100%, 50%, and 10% ethanol, 10min each (30 min total time). After each incubation, tubes are centrifuged at 10,000xG for 2min. Pellets are resuspended in extraction buffer, incubated at 95°C (20min), placed on ice and briefly centrifuged. After 6μl (60μg) RNase is added to each tube, samples are incubated for 30min at RT, and then 8μl (160μg) Proteinase K is added. The suspension (100μl aliquots/well) is transferred to PIXUL plate, sonicated (Pulse=50, PRF=1.0, Burst=20, T=30min), transferred to 0.5mL tubes, boiled for 10min, chilled on ice, and centrifuged at 16,000xG for 15min. Supernatants containing ssDNA are used immediately in Matrix methylated DNA immunoprecipitation MeDIP or can be stored frozen (-20°C) for future use.

#### 96-well plate Matrix-MeDIP-seq

96-well microplate-based Matrix methylated DNA immunoprecipitation (Matrix-MeDIP) of ssDNA was done as previously described where 5mC antibody was attached to wall wells via Protein A (28,42). Well walls were blocked with 5% BSA. PIXUL fragmented ssDNA was used as input. After washes, immunocaptured ssDNA was recovered by incubation with Proteinase K (200ng/µL) in 100µL/well of elution buffer (55°C for 45min, 95°C for 10min, then 4°C). MeDIP was validated with PCR. Sequencing libraries were prepared using xGen ssDNA library kit (IDT). Library validation was performed as described below. Libraries were sequenced using NextSeq 2000.

#### Library validation

After library generation for each method, each library was validated, with Collibri Library Quantification Kit PCR used to verify adapter attachment, organ-specific-primer PCR used to show retained specificity, and Agilent 4200 TapeStation used to assess base pair length and library quality.

#### The metrics for inter-tumor and intra-tumor heterogeneity, the heterolon, *H*

We suggest that variation within and across tissues can be analyzed using a string of elements (bins) having primary (for example, epigenetic mark, *epi*), secondary (gene, *gene*), and tertiary (gene site, *site*) tiers, which we write as epi-gene-site, and suggest calling it the *epi-gene-site* heterolon. The heterolons would have N+1 string elements. We quantitate heterolons across tissue samples with the coefficient of variation, CV (SDEV divided by the mean), calculated for a given heterolon, e.g., *epi-gene-site* CV. Operationally, sample measurements can be grouped into unique heterolons and CVs calculated using data spreadsheets (e.g. Excel).

#### Gene Ontology (GO) analysis

Read counts were generated from the BAM files using <u>featureCounts</u>. <u>Column join</u> was then used to create a single count matrix from the read counts, denoted *countdata*. Annotation data was generated using <u>annotateMyIDs</u> to combine the relevant *countdata* with SYMBOL, GENENAME, and GO columns into a single *annodata* file. Factor data was generated by uploading a user-formatted text file with sample labels to Galaxy, denoted *factordata*. Finally, the *countdata*, *annodata*, and *factordata* files were combined using <u>limma</u> to contrast MGMT methylated and unmethylated samples (limma-voom; contrast of interest: Methylated vs Unmethylated; filter lowly-expressed genes; CPM = 0.5; minimum samples = 2; Output options: Output Normalized Counts Table). The resulting Normalized Counts Table was then exported to Excel. The genomic locations were sorted according to the greatest read count differential (corresponding to greatest differential methylation) between MGMT methylated and unmethylated samples, and the GO ID numbers were matched to specific GO names using the text file found at https://current.geneontology.org/ontology/go-basic.obo.

## SUPPLEMENTAL FIGURES

**Fig. S1.**
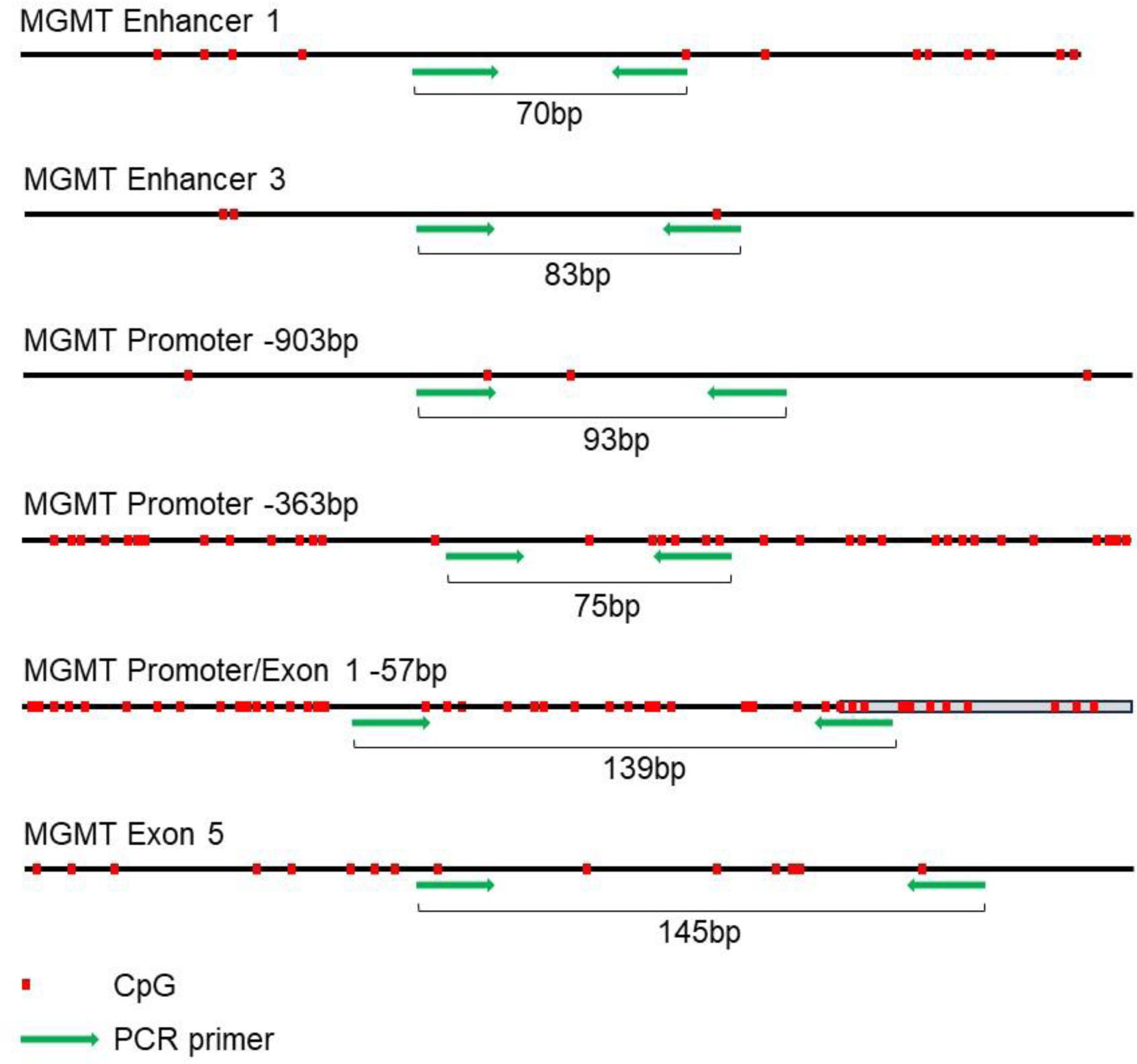
Cartoons showing location of CpG islands and location of PCR amplicons. The average size of the sheared DNA fragments is 200-250bp. As such, the MeDIP qPCR amplicons will detect CpG approximately 100bp flanking the location of the qPCR amplicon.

**Fig. S2.**
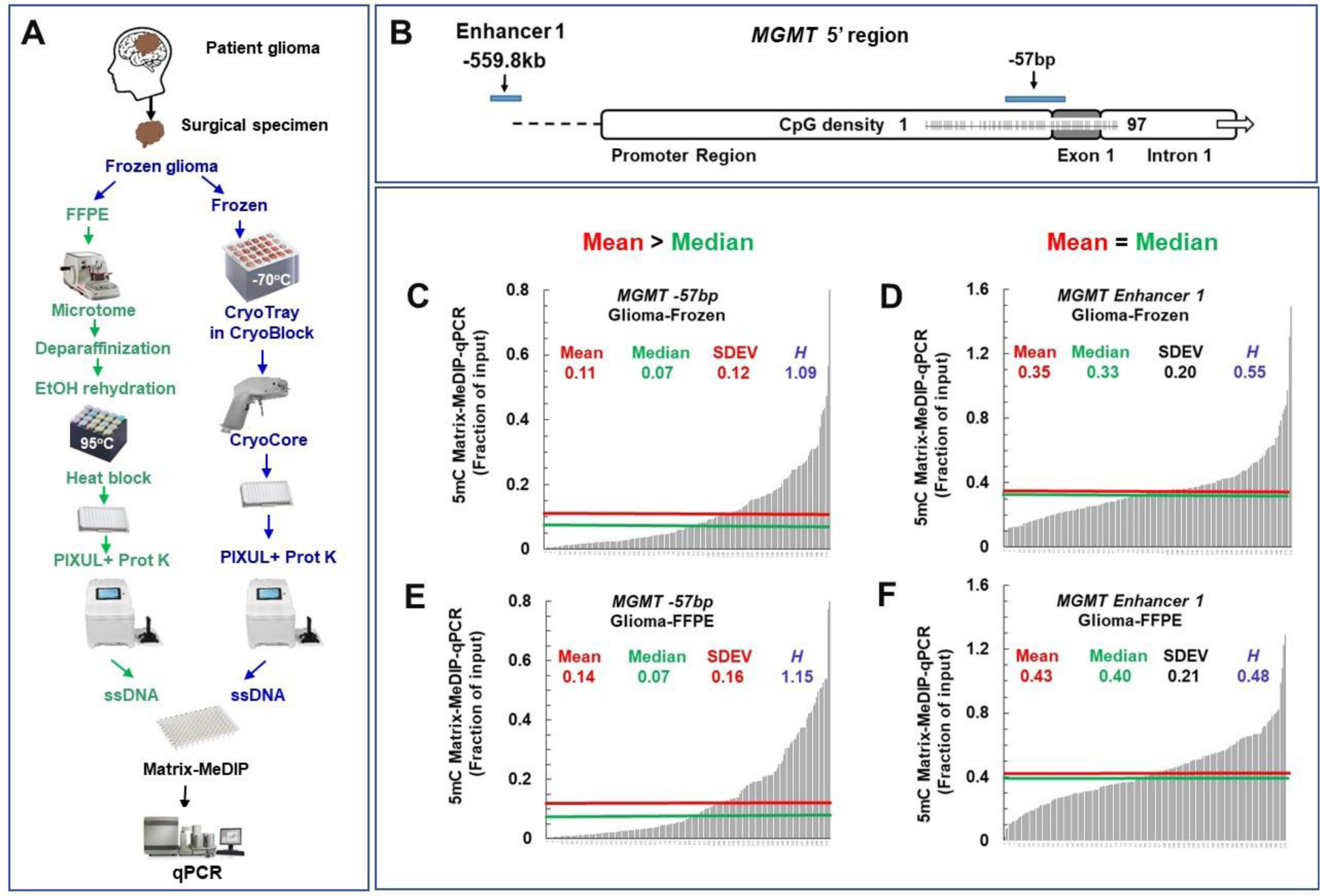
Matrix-MeDIP-qPCR DNA methylation analysis of paired FFPE and frozen gliomas sample collection at *MGMT* promoter and enhancer sites. ***A.*** 96-well plate PIXUL-based protocol was used to isolate DNA from 173 paired FFPE and frozen GBM tissue samples. Matrix-MeDIP was done using 5mC antibody. Briefly, surgical GBM specimen was frozen right after resection. A portion of the frozen specimen was formalin fixed paraffin embedded, FFPE, and slices (5μm) were generated from FFPE blocks using a microtome. After deparaffinization, EtOH rehydration, and heat retrieval, samples were treated in 96-well PIXUL, and DNA was isolated using proteinase K. Frozen GBM tissues were placed in pockets of 24-well CryoTrays and sampled with the CryoCore, cores were treated in 96-well PIXUL, and DNA was isolated using proteinase K. FFPE and frozen samples’ DNA was boiled (ssDNA) and used in Matrix-MeDIP-qPCR using 5mC antibody. ***B.*** *MGMT* 5’ region showing sites of PCR primers. ***C-D.*** Distribution of 5mC signal (expressed as fraction of input) in the 5’ MGMT region; -57bp CpG island site (***C***) and -560kb enhancer site (***D***) from DNA isolated from frozen specimen, ***E-F***. Distribution for FFPE tissues. Shown in each graph are the Mean, Median, SDEV, and *H* (heterolon) values. The red and green lines show the mean and median values respectively.

**Fig. S3.**
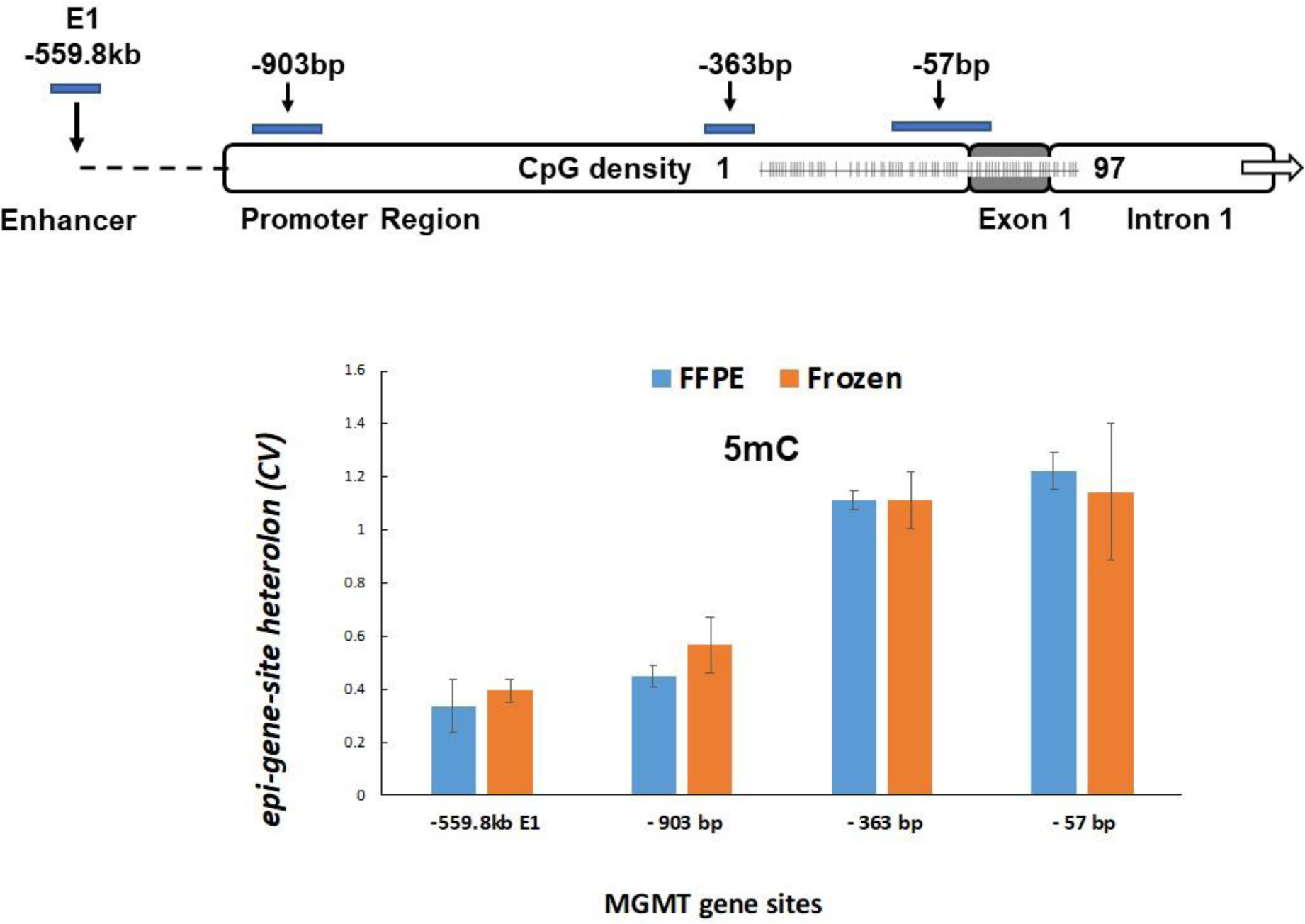
MGMT DNA methylation *epi-gene-site* heterolon CVs (i.e. each bar represents a heterolon) are the same for fresh frozen (FF) and FFPE tumors but are site-dependent. 5mC levels were measured by Matrix ChIP-qPCR at the indicated *MGMT* sites. 5mC CVs were calculated separately for 26 HM and 12 UW FF and FFPE tumor pairs and then averaged for the two sets of samples.

**Fig. S4.**
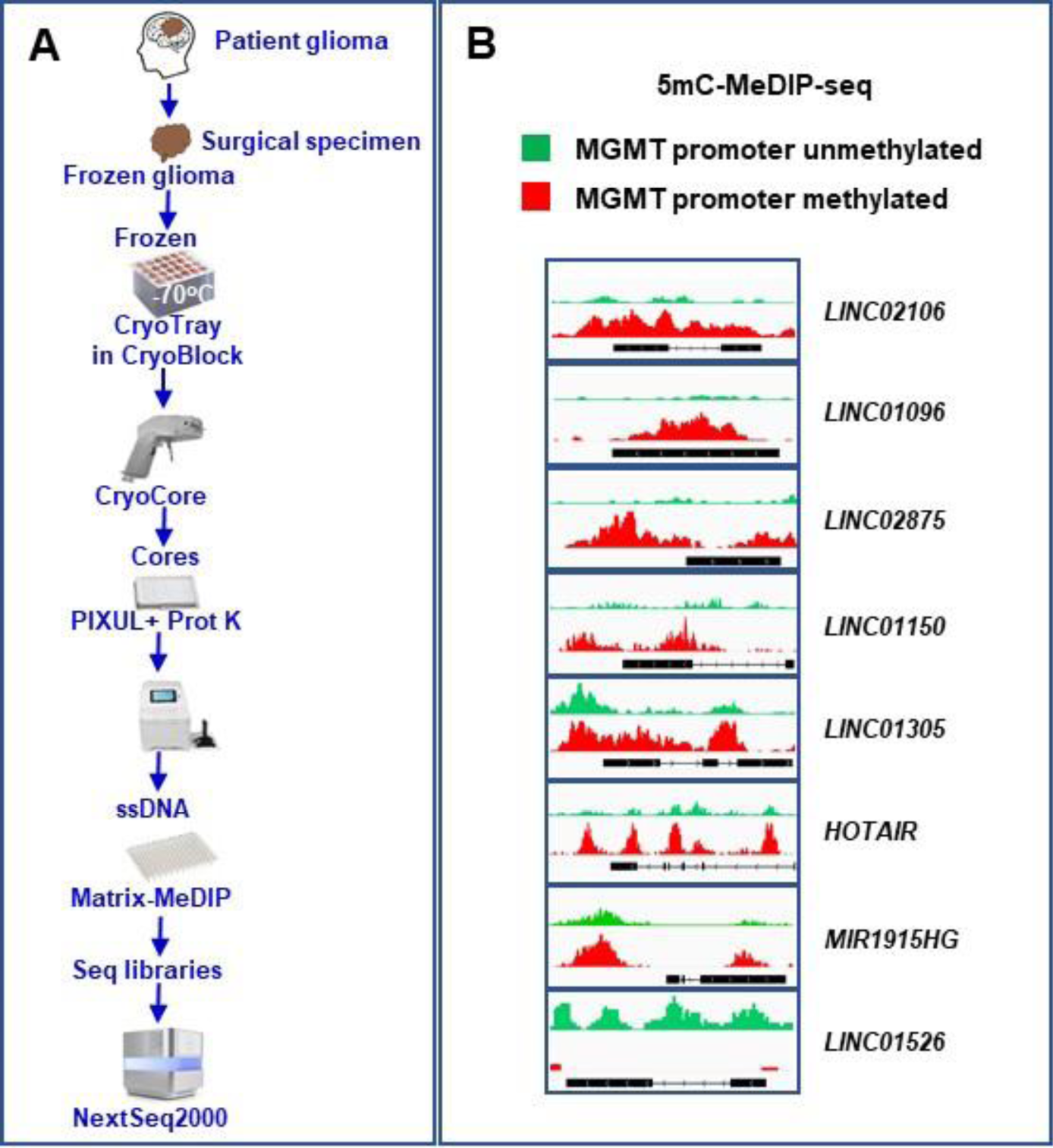
IGV MeDIP-seq snapshot of DNA methylation of long non-protein coding intragenic RNA, lncRNA, in gliomas with methylated (*red*) and unmethylated (*green*) *MGMT* promoter. *A.* Frozen gliomas in CryoTray were sampled with the CryoCore, with cores jetted into two 96-well plates to isolate DNA. ssDNA was used in 5mC Matrix-MeDIP and then to generate sequencing libraries. *B.* IGV snapshots of 5mC MeDIP-seq.

## TABLES

**Table S1.**
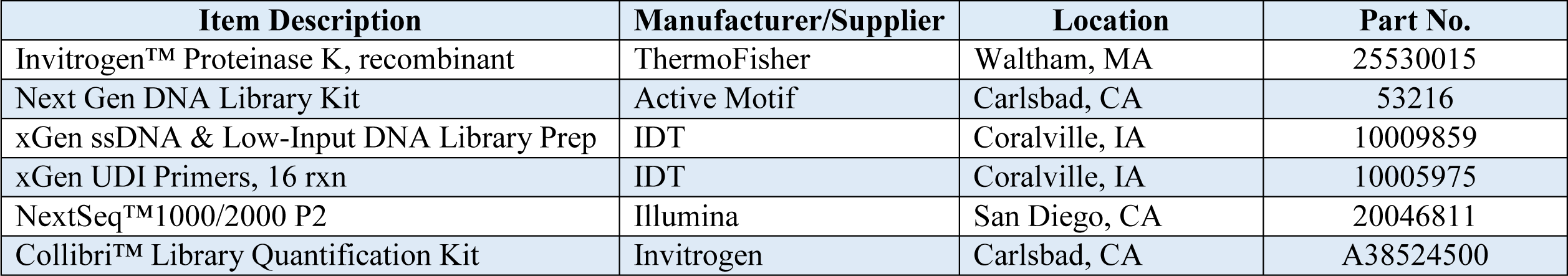
Kits and enzymes.

**Table. S2.**
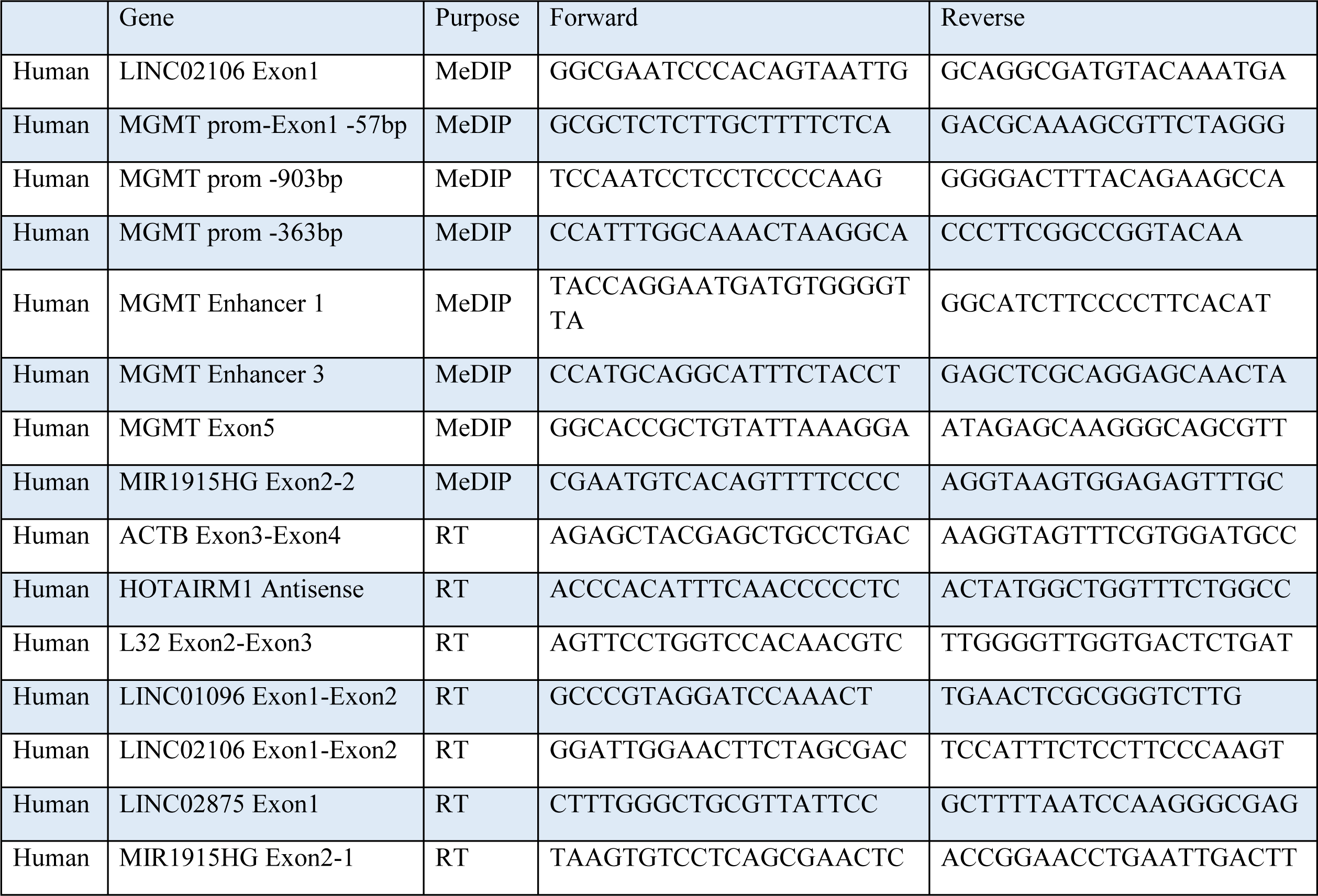
qPCR primers.

**Table. S3.** Antibodies.

| **Antibody** | **Assay** | **Catalog No** | **Source** | **Manufacturer** |
| --- | --- | --- | --- | --- |
| 5mC | MeDIP-seq | AMM99021- | Mouse monoclonal 33D3 | Aviva Systems Biology |
| IgG | ChIP-qPCR/seq | I-2000 | Mouse serum | Vector Laboratories |

